# Revisiting the effect of red on competition in humans

**DOI:** 10.1101/086710

**Authors:** Laura Fortunato, Aaron Clauset

## Abstract

Bright red coloration is a signal of male competitive ability in animal species across a range of taxa, including non-human primates. Does the effect of red on competition extend to humans? A landmark study in evolutionary psychology established such an effect through analysis of data for four combat sports at the 2004 Athens Olympics [1]. Here we show that the results do not replicate in an equivalent, independent dataset for the 2008 Beijing Olympics, and that there is substantial variation in the fraction of wins by red across sports in both years. We uncover a number of shortcomings with the research design, analysis, and interpretation underlying the original results. For example, the variation observed in the data may reflect bias towards wins by one color over the other, linked to specific features of the tournament structure for the sports analysed. Reanalysis of the data to address these shortcomings indicates that there is no evidence for an effect of red on the outcomes of Olympic combat sports. Our results refute past claims based on analysis of this system, challenging the related notion that any effect of red in human competition is an evolved response shaped by sexual selection.

---

In animal species across a range of taxa, bright coloration is a secondary sexual character acting as a signal of male competitive ability [2]. In mandrills, for example, male rank is determined through contest competition, with marked reproductive skew in favor of top-ranking individuals. High rank is associated with better reproductive outcomes also in females, but here rank is inherited from the mother instead. As expected within the framework of Darwinian sexual selection [3, 4], the extent and intensity of red skin on the face of adult individuals vary with rank in males, but not in females [2, 5].

The relationship between red coloration and competition in non-human primates and other taxa raises an intriguing question [1]: does red have an effect on the outcome of human competitive interactions, shaped by similar evolutionary processes? Of course, humans do not present natural displays of conspicuous secondary sexual coloration. However, fluctuations in blood flow to the skin are linked to a range of emotional states, including anger and fear. This response may serve as a subtle cue of relative dominance during aggressive encounters, echoing the sexually selected response to red in other species.

Hill & Barton [1] reasoned that the effect may extend to artificial stimuli, for example wearing red during a physical contest. In an ingenious first test of this hypothesis, they exploited a feature of four Olympic combat sports: boxing, taekwondo, Greco-Roman wrestling, and free-style wrestling. In these sports, contestants compete in pairs as red vs. blue, with distinctively colored clothing and/or equipment. If red confers a competitive advantage, as predicted, then contestants wearing red would be more likely to defeat their opponents, and significantly more than half the contests would end in a win by red.

Using data for the men’s divisions of the four sports at the 2004 Athens Olympics, Hill & Barton [1] reported a series of results broadly in line with the prediction [1] (Fig. 1**a**; Supplementary Information). No effect was found in corresponding data for the women’s divisions (featured only in taekwondo and in free-style wrestling) [6]. These patterns were taken to support the hypothesis of an effect of red in human competitive interactions: red enhances performance, possibly acting as a cue of relative dominance when factors such as skill or strength are equally matched. At the proximate level, the effect may operate through psychological or physiological (e.g., hormonal) influences on the red-wearing competitor, on his opponent, or both [1, 6].

**FIG. 1.**
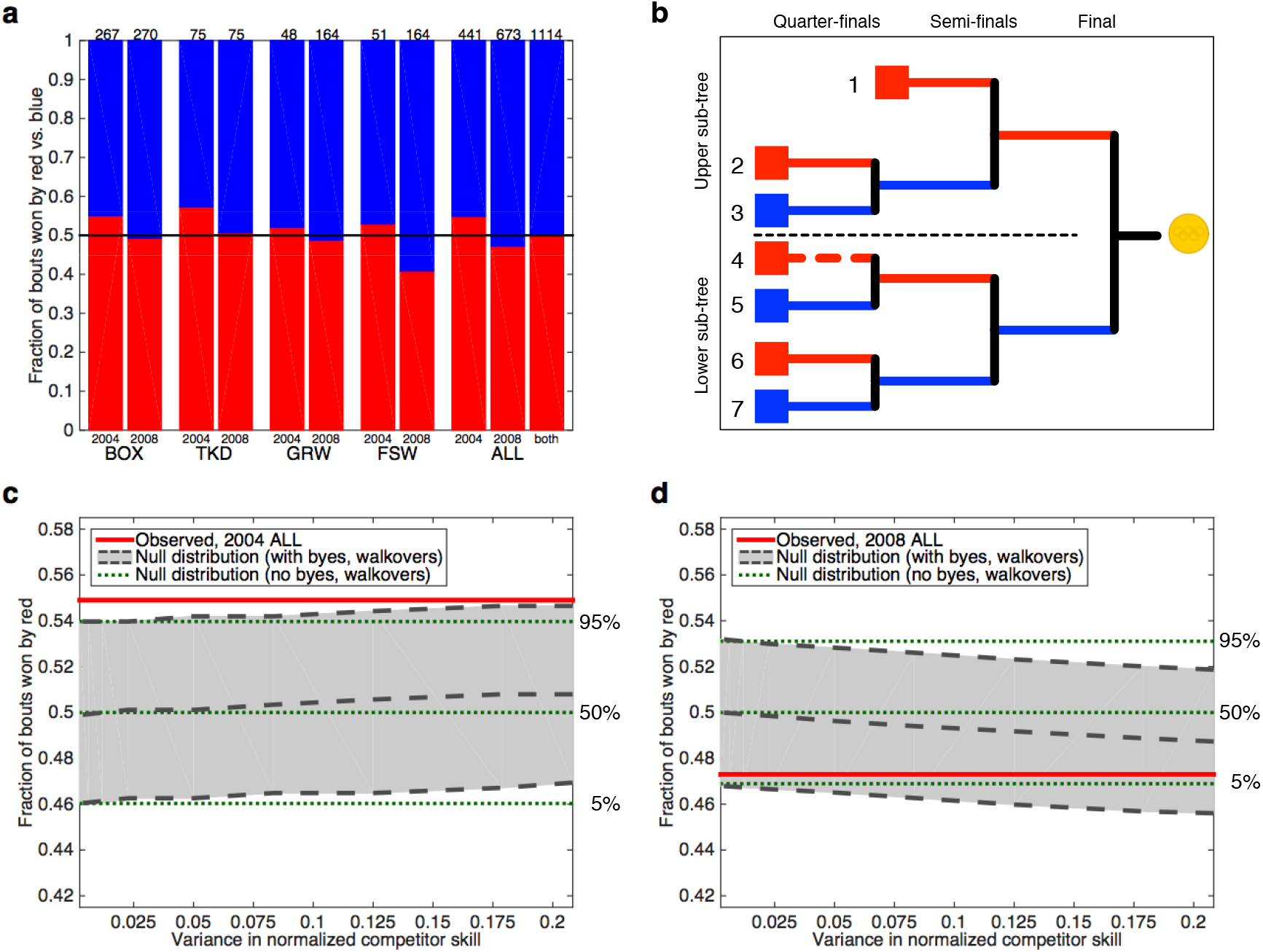
Testing the effect of red in Olympic contests. **(a)** Fractions of bouts won by contestants wearing red vs. blue in the male divisions of boxing (BOX), taekwondo (TKD), Greco-Roman wrestling (GRW), and free-style wrestling (FSW), and aggregated over the four sports (ALL) by year (2004, 2008) and over the two years (both), at the 2004 Athens Olympics and the 2008 Beijing Olympics. The number of bouts in each group is reported above the corresponding bar. The horizontal line shows *f*_red_ = 0.5. **(b)** Schematic representation of the structure of a single-elimination tournament for *n* = 7 contestants. Because *n* is not a power of 2, the outermost round is incomplete. In this case, contestant 1 does not compete in the quarter-final round, i.e., he is byed to the semi-final round. In each contest, or bout, the contestant at the top of the bracket wears red, the one at the bottom wears blue. For example, contestants 2 and 3 wear red and blue, respectively, in the quarter-final round. The winner of this bout proceeds to the semi-final round (in blue), where he faces contestant 1 (in red). The bout between contestants 4 and 5 is won by walkover (dashed red line, indicating that contestant 4 withdrew or failed to show up). Contestant 5 proceeds to the semi-final round (in red), where he faces the winner of the 6–7 bout (in blue). **(c, d)** Quantiles for the distribution of the fraction of wins by red *f*_red_ under the null hypothesis on (i) the observed tournament structures for the two Olympic competitions (dashed grey lines, grey fill), and (ii) equivalent tournaments with no missing bouts (dotted green lines). The solid red line shows the observed value of *f*_red_. For the observed tournaments, the null distribution shifts away from a mean of *f*_red_ = 0.5 as the variance in competitor skill increases, with bias towards wins by red in 2004 and blue in 2008. See text for details.

Hill & Barton’s [1] results have inspired a large body of work in evolutionary psychology and related areas over the past decade, and their influence extends beyond this domain [see reviews in 7–10]. For example, the effect of red on human behavior has come to be regarded as one of the best established in the field color psychology, with diverse practical applications. In this field, Hill & Barton’s [1] study is considered a major theoretical advance [8], which “[d]ocuments parallels between the human and nonhuman response to color” [8, p. 115].

We present multiple lines of analysis, together with new data, to evaluate the foundational finding to this body of work — Hill & Barton’s [1] demonstration of an effect of red on the outcome of Olympic combat sports. We use two independent but fully equivalent datasets: the data provided by Hill & Barton [1] for the 2004 Athens Olympics, and data we obtained separately for the 2008 Beijing Olympics. Both datasets relate to the male divisions of the four sports analysed in [1] (Supplementary Information).

Crucially, Hill & Barton’s [1] results do not replicate in the 2008 data (Supplementary Information). In fact, the fraction of wins by red varies substantially across sports in both years (Fig. 1**a**), being exactly half in the data aggregated over the two years (*f*_red_ = *n*_red_*/n*_tot_ = 560*/*1114 = 0.503). Indeed, the largest bias observed in any sport in either year is towards wins by blue, not red (*f*_red_ = 67*/*164 = 0.409 in 2008 free-style wrestling).

This variation in the data invites caution in drawing inferences about the effect of color on the outcomes of Olympic combat sports. It also raises concerns about the validity of Hill & Barton’s [1] results. In re-deriving the analytical approach used in [1], we uncovered several shortcomings with the underlying research design, analysis, and interpretation (Supplementary Information). A key issue is that the pattern observed in the data (Fig. 1**a**) is not simply the product of random variation. Rather, it may also reflect bias towards wins by one color over the other, linked to specific features of the tournament structure for the sports analysed. We investigated this issue mathematically and numerically (Methods), and we discuss related insights here because they have general relevance, beyond the system under study. Several other issues that apply specifically to the research design, analysis, and interpretation in [1] are detailed in the Supplementary Information.

In boxing, taekwondo, and wrestling, the competition for a given weight class is arranged as a single-elimination tournament (Fig. 1**b**). Generally, the winner of a contest, or bout, proceeds to the next round in the competition tree. In boxing and in wrestling, in each bout the contestant at the top of the bracket wears red, the one at the bottom wears blue; the assignment is reversed in taekwondo. A contestant’s relative position, and thus the color he wears, may change between bouts, as he progresses through rounds in the tournament (Supplementary Information).

The assigment procedure does not, in itself, introduce bias towards wins by one color over the other. It can do so, however, if coupled with variance in skill among the contestants, together with incompleteness in the tournament tree. Two sources of incompleteness are byes and walkovers (Fig. 1**b**). Both result in missing bouts, but whereas walkovers can occur anywhere on the tree, byes are always placed in the outermost round. In particular, byes are “stacked” in the upper sub-tree or in the lower one (Supplementary Information).

Byes and walkovers can both result in bias. The underlying mechanism is asymmetric selection on contestant skill in the two sub-trees feeding into a bout (Supplementary Information). In the case of byes the bias is systematic: as we show in the Supplementary Information, direction and magnitude can be predicted based on a few key factors. For example, if the assignment procedure is as in boxing and in wrestling (i.e., in each bout the contestant at the top of the bracket wears red), then the null distribution for the fraction of wins by red is centered at *f*_red_ *>* 0.5 with byes stacked in the lower sub-tree; it is centred at *f*_red_ *<* 0.5 with byes stacked in the upper sub-tree. The magnitude of the bias depends on the number of byes relative to the total number of bouts in the outermost round, and it is greater for odd numbers of byes than for adjacent even numbers of byes. In the case of walkovers any bias is idiosyncratic, in the sense that direction and magnitude depend on the specific configuration.

If a competition tree presents both systematic and idiosyncratic biases, due to byes and walkovers respectively, these may interact synergistically or antagonistically — their combined effect also depends on the specific configuration. Additionally, in taekwondo and in wrestling the combined effect arises from interaction of any bias in the main tournament with any bias in the consolation round(s) (Supplementary Information).

In all of the main tournaments analysed here, the position of contestants on the outermost round of the tree was determined at random (i.e., there was no seeding by skill; Supplementary Information). Consequently, which colors a contestant wore at the outset of a tournament was independent of his skill. This excludes the possibility that any observed bias arose from the allocation of strong contestants to specific positions on the tree — a mechanism posited for single-elimination tournaments featuring seeding by skill in judo [11].

To determine whether bias towards wins by one color over the other applies to the 2004 and 2008 datasets, we numerically calculated the distribution of wins by red under the null hypothesis (no effect of red) for (i) the observed tournaments, and (ii) equivalent tournaments with no missing bouts. The null distribution for the observed tournaments shifts towards wins by one color as the variance in competitor skill increases — specifically, towards red in 2004 and towards blue in 2008 (Fig. 1**c, d**). As discussed in the Supplementary Information, the shift is driven mainly by Greco-Roman and free-style wrestling. Compared to the other sports, wrestling included a large number of missing bouts relative to the size of the tournaments. Crucially, following substantial changes in tournament structure implemented between the two competitions, byes were stacked in the lower subtree in 2004, and in the upper one in 2008. This pattern is consistent with null distributions centered at *f*_red_ *>* 0.5 and *f*_red_ *<* 0.5, respectively, for contestants in the top bout position wearing red (Supplementary Information). These findings indicate that the results in [1] likely overstate the evidence for an effect of red in the 2004 data. In particular, for data relating to wrestling, a correctly parameterized test of Hill & Barton’s [1] hypothesis cannot be constructed without knowledge of the variance in competitor skill. Absent this information, a conservative approach is to exclude the corresponding data from further analysis.

Even at the highest levels of variance in skill, the overall bias is minimal in boxing and in taekwondo (Supplementary Information). Therefore, the corresponding data can be used to test Hill & Barton’s [1] hypothesis. For comparison with the analysis in [1], we conducted nine binomial tests (*H*_0_ : *f*_red_ ≤ 0.5; *H*_A_ : *f*_red_ *>* 0.5; *α* = 0.05), relating to the fraction of bouts won by red in boxing and in taekwondo, individually by year and aggregated over all sport/year combinations. We adjusted for multiple hypothesis testing using four methods, controlling both the family-wise error rate [12–14] and the false discovery rate [15, 16] (Supplementary Information).

Evaluated as a whole, the results return no evidence for an effect of red (Table I). Even based on the raw *p*-values, the null hypothesis is rejected in only one of the nine tests. Inspection of the confidence intervals and of the adjusted *p*-values suggests that this is likely a case of spurious statistical significance (Supplementary Information).

**TABLE I.**
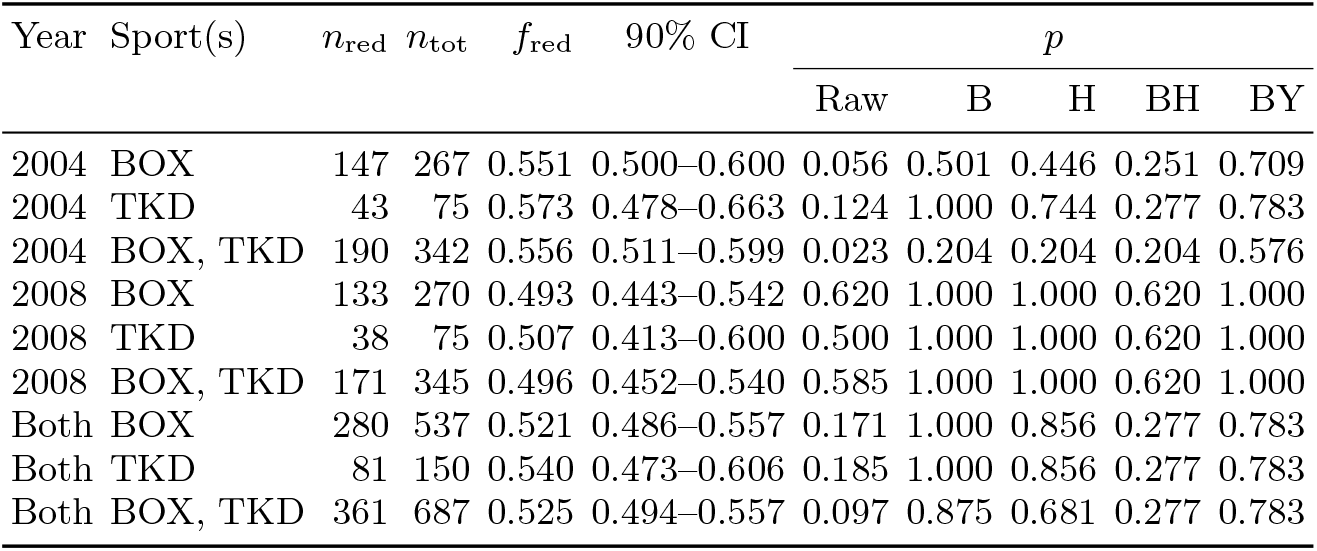
Results of one-sided binomial tests in data for boxing (BOX), taekwondo (TKD), and aggregated over the two sports (BOX, TKD), at the 2004 Athens Olympics and the 2008 Beijing Olympics. The tests compare the number of bouts won by red, *n*_red_, to the total number of bouts, *n*_tot_, with *f*_red_ = *n*_red_*/n*_tot_. Included are the raw *p*-values (*H*_0_ : *f*_red_ ≤ 0.5; *H*_A_ : *f*_red_ *>* 0.5; *α* = 0.05), the 90% confidence intervals for *f*_red_, and the *p*-values adjusted for multiple hypothesis testing using four methods (B: Bonferroni [12, 13]; H: Holm [14]; BH: Benjamini & Hochberg [15]; BY: Benjamini & Yekutieli [16]).

We have presented the most extensive evaluation to date of Hill & Barton’s [1] hypothesis of an effect of red on the outcomes of Olympic combat sports. Our findings refute claims about the role of color in human competitive interactions, based specifically on analysis of this system [1]. They also suggest that caution is required in interpreting related results derived from analysis of other systems [e.g., competition outcomes in other sports; reviewed in 7–10]. Our insights do not apply directly to experimental studies investigating the effect of red on competition, or on other aspects of human behavior [also reviewed in 7–10]. However, they do undermine the widespread reliance of these studies on Hill & Barton’s [1] results as evidence that any such effect has an evolutionary basis (Supplementary Information).

More broadly, our work challenges Hill & Barton’s [1] conjecture of parallels between a sexually selected response to red in humans, mandrills, and other species. In what way evolution has shaped the human response to color, and how this is reflected in present-day human behavior, remain open questions.

## Methods

Details of the data collection and analysis are in the Supplementary Information. The distributions of *f*_red_ under the null hypothesis, shown in Fig. 1**c, d**, were obtained by Monte Carlo simulation of competition on individual single-elimination tournaments. Results were then aggregated for analysis. Contestants were assigned randomly to initial tournament positions, with skill levels drawn i.i.d. from a symmetric Beta distribution *x* ∼ Beta(*β, β*). Bout outcomes were evaluated progressively over rounds. When a pair of competitors *r* and *b* faced off, *r* advanced to the next round with probability *x*_*r*_*/*(*x*_*r*_ + *x*_*b*_) [17] (Supplementary Information)

## Notes

### Competing Interest Statement

The authors have declared no competing interest.

## References

[1] Hill, R. A. & Barton, R. A. 2005 Red enhances human performance in contests. Nature 435, 293–293.

[2] Setchell, J. M. 2015 Color in competition contexts in nonhuman animals. In Handbook of color psychology (edited by A. J. Elliot, M. D. Fairchild, & A. Franklin), chap. 26, pp. 546–567. Cambridge: Cambridge University Press.

[3] Darwin, C. 1871 The descent of man, and selection in relation to sex. London: John Murray.

[4] Darwin, C. 1876 Sexual selection in relation to monkeys. Nature 15, 18–19.

[5] Setchell, J. M. 2016 Sexual selection and the differences between the sexes in mandrills (Mandrillus sphinx). American Journal of Physical Anthropology 159, 105–129.

[6] Barton, R. A. & Hill, R. A. 2005 Sporting contests: Seeing red? Putting sportswear in context (reply). Nature 437, E10–E11.

[7] Wiedemann, D., Barton, R. A., & Hill, R. A. 2012 Evolutionary perspectives on sport and competition. In Applied evolutionary psychology (edited by S. C. Roberts), chap. 18, pp. 290–307. Oxford: Oxford University Press.

[8] Elliot, A. J. & Maier, M. A. 2014 Color psychology: effects of perceiving color on psychological functioning in humans. Annual Review of Psychology 65, 95–120.

[9] Maier, M. A., Hill, R. A., Elliot, A. J., & Barton, R. A. 2015 Color in achievement contexts in humans. In Handbook of color psychology (edited by A. J. Elliot, M. D. Fairchild, & A. Franklin), chap. 27, pp. 568–584. Cambridge: Cambridge University Press.

[10] Rowland, H. M. & Burriss, R. P. 2017 Human colour in mate choice and competition. Philosophical Transactions of the Royal Society of London B: Biological Sciences 372. http://dx.doi.org/10.1098/rstb.2016.0350.

[11] Dijkstra, P. D. & Preenen, P. T. 2008 No effect of blue on winning contests in judo. Proceedings of the Royal Society of London B: Biological Sciences 275, 1157–1162.

[12] Dunn, O. J. 1959 Estimation of the medians for dependent variables. Annals of Mathematical Statistics 30, 192–197.

[13] Dunn, O. J. 1961 Multiple comparisons among means. Journal of the American Statistical Association 56, 52–64.

[14] Holm, S. 1979 A simple sequentially rejective multiple test procedure. Scandinavian Journal of Statistics 6, 65–70.

[15] Benjamini, Y. & Hochberg, Y. 1995 Controlling the false discovery rate: a practical and powerful approach to multiple testing. Journal of the Royal Statistical Society: Series B 57, 289–300.

[16] Benjamini, Y. & Yekutieli, D. 2001 The control of the false discovery rate in multiple testing under dependency. The Annals of Statistics 29, 1165–1188.

[17] Bradley, R. & Terry, M. 1952 Rank analysis of incomplete block designs. i. The method of paired comparisons. Biometrika 39, 324–345.

