## Supplementary Information File for "Revisiting the effect of red on competition in humans"

### Revisiting the effect of red on competition in humans (Supplementary Information)

**S7 Discussion** 52

**References** 54

**Acknowledgments** 56

**Session information** 57

### S1 Outline

In Section S2 we present descriptions of relevant aspects of the competitions for the four sports analyzed by Hill & Barton [1], relating to the 2004 Athens Olympics and the 2008 Beijing Olympics.

In Section S3 we provide information on acquisition and processing of the data for analysis, together with descriptive statistics for each sport/year combination.

In Section S4 we replicate the results reported in [1], and we devise an alternative analytical approach to address key shortcomings of the original one.

In Section S5 we investigate the mechanism leading to bias towards wins by one color over the other in single-elimination tournaments.

In Section S6 we triangulate insights from the different lines of analysis into a final set of tests of the prediction of an effect of red on the outcome of Olympic combat sports.

In Section S7 we conclude with an overall summary of our findings, focusing on their implications for the hypothesis of an effect of red on human competition, and on human behavior more generally, in the context of the substantial body of work that has developed building on Hill & Barton's [1] influential study.

Readers interested specifically in the discussion presented in the main text can skip directly to the extensive summaries in Sections S2.5, S4.4 and S5.4, followed by Sections S6 and S7.

#### S2 Structure of the sports analysed

Our aim in this section is three-fold. First, we seek to provide readers with a mechanistic understanding of the competition structure for the different sports, focusing on specific features leading to bias towards wins by one color over the other in the outcomes of the competitions. Second, we highlight the equivalence between the 2008 data presented here and the 2004 data analysed by Hill & Barton [1]. Third, we outline changes in the competition structure introduced at the 2012 London Olympics in two of the four sports, which prevent extension of the analysis to data relating to this competition.

Below we define terminology that applies to single-elimination tournaments generally, before describing specific features of the tournament structure and related aspects for the different sports (male divisions) — including the color assignment procedure, the placement of winners, and a comparison across competition years.

##### S2.1 Terminology

In the four sports analysed by Hill & Barton [1] (male divisions), the competition for a given weight class was arranged as a single-elimination tournament (also known as "knock-out" or "sudden death"). As illustrated in Fig. 1**b** in the main text, in this type of tournament contestants compete in pairs. The winner of a contest, or bout, proceeds to the following round in the competition tree. A contestant's relative position in a given bout (top or bottom of the bracket) determines the color he wears in that bout. His relative position may change between bouts, as he progresses through rounds in the tournament.

Two possible sources of incompleteness in a single-elimination tournament are byes and walkovers, also illustrated in Fig. 1**b** in the main text. Byes are used if there are fewer than the number of contestants required to fill the outermost round in the tree; in this case, byed contestants effectively bypass the outermost round, "skipping" to the following round. A walkover involves a contestant winning the bout by default, because his opponent forfeited the contest (e.g., by withdrawing or by failing to show up).

Both byes and walkovers result in missing bouts, but whereas walkovers can occur anywhere on the tree, byes are always placed in the outermost round (Fig. 1**b** in the main text). In particular, byes can be drawn from the highest position in the round going down, or from the lowest position going up. These procedures result in byes "stacked" in the upper vs. lower sub-trees, respectively, in the sense that there will be more byes in upper sub-tree in one case, and more in the lower sub-tree in the other.

For a tournament with $n \geq 2$ contestants, there are $n_{\text{rounds}} = \lceil \log_2 n \rceil$ rounds in the competition tree. The number of contestants required to fill the outermost round is $2^{n_{\text{rounds}}}$, resulting in $n_{\text{byes}} = 2^{n_{\text{rounds}}} - n$ byes and $n^{\text{out}}_{\text{bouts}} = (n - n_{\text{byes}})/2$ bouts in this round. Qualitatively, the implication is that if the number of contestants in the tournament is not a power of 2, then there will be one or more byes in the outermost round. The contestants who are not byed compete in this round, with the losers eliminated from the tournament.

As a result, the number of contestants in the following round is reduced to a power of 2. This leads to a series of complete rounds after an incomplete outermost one.

The number of bouts across all rounds is thus $n^{\text{tot}}_{\text{bouts}} = n^{\text{out}}_{\text{bouts}} + 2^{n_{\text{rounds}}-1} - 1$, which simplifies to $n^{\text{tot}}_{\text{bouts}} = n - 1$. The intuition here is that, except for the overall winner, every contestant in the tournament will lose exactly one bout. Hence the total number of bouts on the tree (corresponding to its internal nodes) is one less than the initial number of contestants (corresponding to its leaves, or tips).

We define the completeness fraction linked to byes as $\rho = (n - n_{\text{byes}})/2^{n_{\text{rounds}}}$, with $\rho = 1.00$ for a tournament with no byes and $\rho < 1.00$ for a tournament with byes. The average value of $\rho$ across weight classes for a sport in a given year provides a simple measure of how systematically (in)complete the tournaments tended to be.

#### S2.2 Boxing

The male division at the 2004 and 2008 Olympics included 11 weight classes.

##### S2.2.1 Tournament structure

For each weight class, the competition was arranged as a single-elimination tournament. In the 2004 and 2008 tournaments, the number of contestants $n$ ranged from 16 to 29 across weight classes. Tournaments with $n > 16$ had an incomplete outermost round, with $n$ ranging from 27 to 29. Thus, there were byes in the outermost round of these tournaments (Section S2.1). Specifically, some of the contestants received a bye to the round of 16 (eighth-finals), whereas the others competed in the round of 32, with the winners of bouts in this round proceeding to the round of 16. The average completeness fraction linked to byes was $\rho = 0.79$ for the 2004 tournaments and $\rho = 0.79$ for the 2008 tournaments (Section S2.1). The rounds of 16 and of 32 were both referred to as "preliminaries".

In the 2004 and 2008 tournaments, byes were drawn from the highest position in the outermost round going down, resulting in byes stacked in the upper sub-tree. The initial position of contestants on the tree (i.e., the seeding of the outermost round) was drawn by manual lot, and thus at random; byes were determined through this draw, and thus also at random (Official Report of the XXVIII Olympiad 2: the Games, pag. 277; Sébastien Gillot, pers. comm. Nov. 2013; Janusz Majcher, pers. comm. Nov. 2013).

##### S2.2.2 Color assignment procedure

In the 2004 and 2008 competitions, contestants wore blue or red uniforms (vest and shorts) with matching equipment (headguard, gloves).

In each bout, the contestant at the top of the bracket wore red, the one at the bottom wore blue. Colors were assigned following this procedure throughout the tournament — initially based on the position of contestants in the outermost round, then re-assigned accordingly as contestants proceeded from one round to the next.

##### S2.2.3 Placement of winners

The winner of the bout in the final round got first place (gold), the loser second place (silver). The losers of the two bouts in the semi-finals shared third place (bronze). The losers of the four bouts in the quarter-finals shared fifth place.

##### S2.2.4 Comparison across years

In boxing there were no major structural changes between the 2004 and 2008 competitions. Consequently, Hill & Barton's [1] approach can be readily extended to data relating to the 2008 competition.

Starting with the 2012 London Olympics, the draw used to determine the initial position of contestants on the tree was seeded based on the International Boxing Association (AIBA) ranking and on performance in the World Series of Boxing (WSB) (Sébastien Gillot, pers. comm. Nov. 2013; Janusz Majcher, pers. comm. Nov. 2013), with byes and seeded entries evenly distributed across the tree (see e.g., Appendix E of the AIBA Technical & Competition Rules effective from March 24, 2011).

#### S2.3 Taekwondo

The male division at the 2004 and 2008 Olympics included four weight classes.

##### S2.3.1 Tournament structure

For each weight class, the competition was arranged as a single-elimination tournament with $n = 16$ contestants. Thus, the outermost round was always the round of 16 (eighth-finals), and it was always complete, so there were no byes (Section S2.1). The average completeness fraction linked to byes was thus $\rho = 1.00$ for both the 2004 and 2008 tournaments (Section S2.1).

In the 2004 and 2008 tournaments, the initial position of contestants on the tree was drawn by lot, hence the pairing of contestants was by random selection, without consideration of skill (Jeongkang Seo, pers. comm. Oct.–Nov. 2013).

For comparison, it is useful to note that there were 15 contestants per weight class in the female division of 2004; the one who picked number 1 in the draw was byed and proceeded to the quarter-finals, without a match in the eighth-finals (Jeongkang Seo, pers. comm. Oct.–Nov. 2013).

##### S2.3.2 Color assignment procedure

In the 2004 and 2008 competitions, contestants wore a white uniform (called "dobok", consisting of pants and a long-sleeved top), with blue or red equipment (trunk and head protectors).

In each bout, the contestant at the top of the bracket wore blue, the one at the bottom wore red. Colors were assigned following this procedure throughout the tournament — initially based on the position of contestants in the outermost round, then re-assigned accordingly as contestants proceeded from one round to the next.

##### S2.3.3 Placement of winners

The winner of the bout in the final round got first place (gold), the loser second place (silver). Repechage was used to determine the ranking of other contestants, with different types used in the 2004 and 2008 competitions.

**2004 Athens Olympics** The upper half of the main tournament tree was referred to as "Pool A", the lower half as "Pool B". There was one repechage contest with six contestants, arranged in the following configuration ("b": blue; "r": red):

```
1 (b) _____
           |__
2 (b) __   |  |
        |__|  |
3 (r) __|     |
              |__ bronze
4 (b) __      |
        |_    |
5 (r) __| |   |
          |___|
6 (r) ____|
```

1 : loser in semi-final round for Pool A
2 : loser in quarter-final round against finalist for Pool B
3 : loser in eighth-final round against finalist for Pool B

4 : loser in eighth-final round against finalist for Pool A
5 : loser in quarter-final round against finalist for Pool A
6 : loser in semi-final round for Pool B

The winner of the bout in the bronze round was awarded third place (bronze).

Note that the repechage contest was "symmetric" in the allocation of color, in the sense that losers of bouts in the semi-final round wore blue in the top branch (1) and red in the bottom one (6); losers of bouts in the quarter-final round wore blue in the top one (2) and red in the bottom one (5); losers of bouts in the eighth-final round wore red in the top one (3) and blue in the bottom one (4).

**2008 Beijing Olympics** The upper half of the main tournament tree was referred to as "Pool A", the lower half as "Pool B". There were two separate repechage contests, each including three contestants, arranged in the following configuration ("b": blue; "r": red):

```
1 (b) _____
           |__ bronze
2 (b) __   |
        |__|
3 (r) __|

4 (b) __
        |_
5 (r) __| |
          |__ bronze
6 (r) ____|
```

1 : loser in semi-final round for Pool A
2 : loser in quarter-final round against finalist for Pool B
3 : loser in eighth-final round against finalist for Pool B

4 : loser in eighth-final round against finalist for Pool A
5 : loser in quarter-final round against finalist for Pool A
6 : loser in semi-final round for Pool B

The winners of bouts in the bronze rounds shared third place (bronze), the losers got fifth place, and the other two contestants got seventh place.

Note that the two repechage contests were "symmetric" in the allocation of color, in the sense that losers of bouts in the semi-final round wore blue in the top contest (1) and red in the bottom one (6); losers of bouts in the quarter-final round wore blue in the top one (2) and red in the bottom one (5); losers of bouts in the eighth-final round wore red in the top one (3) and blue in the bottom one (4).

##### S2.3.4 Comparison across years

In taekwondo there were structural changes between the 2004 and 2008 competitions, affecting specifically the repechage rounds. However, as discussed in Section S2.3.3, the structure of the repechage contests was "symmetric" in the allocation of color in both the 2004 and 2008 competitions. Consequently, these changes do not invalidate extension of Hill & Barton's [1] approach to data relating to the 2008 competition.

Starting with the 2012 London Olympics, the World Taekwondo Federation (WTF) introduced seeding of the outermost round of the tournament based on world rankings (Jeongkang Seo, pers. comm. Oct.–Nov. 2013).

##### S2.4 Wrestling

The male division at the 2004 and 2008 Olympics included seven weight classes in each Greco-Roman and free-style wrestling.

Broadly, the two sports differ in forbidding (Greco-Roman wrestling) vs. allowing (free-style wrestling) holds and attacks below the waist. They do not differ in the aspects outlined below.

###### S2.4.1 Tournament structure

The tournament structure changed substantially between the 2004 and 2008 competitions.

**2004 Athens Olympics** The contestants in each weight class initially competed in a series of randomly determined elimination pools, with each contestant competing against all others in the pool. The contestant who scored the greatest number of technical points in each pool proceeded to the round of 8 (quarter-finals); classification points were used to break ties (Tony Black, pers. comm. July 2013). The round of 8 was referred to as the "qualification round".

Elimination pools included three or four contestants. Contestants in a pool of three competed in two rounds, those in a pool of four competed in three rounds. The ordering of the pools determined the order in which contestants entered the qualification round, e.g., the winner of Pool 1 entered the qualification round in the highest position in the round, and so on. In this ordering, pools of three preceded pools of four.

In the 2004 competition, the number of contestants in the elimination pools ranged from 19 to 22 across weight classes in the two sports. Contestants were arranged in one of the following configurations:

| | |
|---|---|
| 19 : | 6 elimination pools (5 pools of 3, 1 pool of 4) |
| 20 : | 6 elimination pools (4 pools of 3, 2 pools of 4) |
| 21 : | 7 elimination pools (7 pools of 3) |
| 22 : | 7 elimination pools (6 pools of 3, 1 pool of 4) |

It is important to distinguish between the number of contestants in the elimination pools and the number entering the qualification round from these pools — the latter corresponds to the number contestants in the main competition tree (Section S2.1). Therefore, in the 2004 tournaments the number of contestants $n$ was either six or seven across weight classes in the two sports.

Thus, there were byes in the qualification round of each tournament (Section S2.1). Irrespective of the specific configuration of the elimination pools, six contestants were arranged in two bouts with two byes, seven contestants in three bouts with one bye. Specifically, some of the contestants received a bye to the round of 4 (semi-finals), the others competed in the round of 8, with the winners of bouts in this round proceeding to the round of 4. The average completeness fraction linked to byes was $\rho = 0.61$ for the Greco-Roman wrestling tournaments and $\rho = 0.68$ for the free-style wrestling tournaments (Section S2.1).

Byes were drawn from the lowest position in the qualification round going up, resulting in byes stacked in the lower sub-tree. Given the ordering of the elimination pools outlined above, contestants from pools of four were always byed to the semi-finals; contestants from pools of three may also be byed, depending on the specific configuration. Because the initial placement of contestants into pools was based on a random draw (Tony Black, pers. comm. July 2013), byes were effectively determined through this draw, and thus also at random.

The winners of bouts in the qualification round and byed contestants proceeded to the semi-finals, the losers to the 5–6 final (in configurations with three bouts in the qualification round, these were the two losers with the most points). The winners of the two bouts in the semi-finals proceeded to the 1–2 final, the losers to the 3–4 final. The placement of winners is discussed below.

**2008 Beijing Olympics** For each weight class, the competition was arranged as a single-elimination tournament. In the 2008 tournaments, the number of contestants $n$ ranged from 19 to 21 across weight classes in the two sports. Therefore, each tournament had an incomplete outermost round, and this was always the round of 32. Thus, there were byes in the outermost round of each tournament (Section S2.1). Specifically, some of the contestants received a bye to the round of 16 (eighth-finals), whereas the others competed in the round of 32, with the winners of bouts in this round proceeding to the round of 16. The average completeness fraction linked to byes was $\rho = 0.24$ for the Greco-Roman wrestling tournaments and $\rho = 0.24$ for the free-style wrestling tournaments (Section S2.1). The round of 32 was referred to as "qualifications".

Byes were drawn from the highest position in the outermost round going down, resulting in byes stacked in the upper sub-tree. The initial placement of contestants on the tree, and thus their pairing, was drawn at random; byes were determined through this draw, and thus also at random [see e.g., Articles 8, 12, 14 of the FILA International Wrestling Rules, release Dec. 2006; confirmed in two later versions (updated Feb. 2010 and July 2014), which implies that this approach was used also in 2008].

##### S2.4.2 Color assignment procedure

In the 2004 and 2008 competitions, contestants wore a blue or red wrestling singlet (a one-piece, tight-fitting uniform).

In each bout, the contestant at the top of the bracket wore red, the one at the bottom wore blue. Colors were assigned following this procedure throughout the tournament — initially based on the position of contestants in the outermost round, then re-assigned accordingly as contestants proceeded from one round to the next.

In the 2004 competition, the same assignment procedure applied to bouts in the elimination pools, but the ordering of contestants in a bout based on draw number "switched" between the three rounds of each pool. Specifically, the contestant with the lower draw number in the pair competed at the top of the bracket in Round 1, at the bottom in Round 2, and at the top in Round 3.

##### S2.4.3 Placement of winners

The winner of the bout in the final round got first place (gold), the loser second place (silver). Different procedures were used in the 2004 and 2008 competitions to determine the ranking of other contestants.

**2004 Athens Olympics** The losers of the two bouts in the semi-finals competed for third place (bronze) in the 3–4 final, with the loser of this bout placed fourth. Two losers of the qualification round competed for fifth and sixth place in the 5–6 final, with any other losers from the qualification round placed seventh, and further placements based on the number of classification points scored in the elimination pools. Note that 5–6 finalists often chose not to compete as they could not medal; in these cases, the bout was won by walkover (Tony Black, pers. comm. July 2013).

**2008 Beijing Olympics** Repechage was used to determine the ranking of contestants from third place onwards.

The upper half of the main tournament tree was referred to as the "upper branch", the lower half as the "lower branch". There were two separate repechage contests, each including up to four contestants (depending on the number of contestants, and their placement, in the qualifications round), arranged in the following configuration ("b": blue; "r": red):

```
1 (r) __
        |__
2 (b) __|  |
           |__
   3 (b) __|  |
              |__ bronze
      4 (b) __|

5 (r) __
        |__
6 (b) __|  |
           |__
   7 (b) __|  |
              |__ bronze
      8 (b) __|
```

1 : loser in qualifications round against finalist for upper branch
2 : loser in eighth-final round against finalist for upper branch
3 : loser in quarter-final round against finalist for upper branch
4 : loser in semi-final round against finalist for upper branch

5 : loser in qualifications round against finalist for lower branch
6 : loser in eighth-final round against finalist for lower branch
7 : loser in quarter-final round against finalist for lower branch
8 : loser in semi-final round against finalist for lower branch

The winners of bouts in the bronze rounds shared third place (bronze), the losers got fifth place. Further placements (seventh onwards) were based on the number of classification points scored throughout the tournament (Tony Black, pers. comm. July 2013).

In the 2008 tournaments, no contestants in the upper branch competed in qualifications rounds. Consequently, in no tournament was there a contestant 1, and contestant 2 was therefore byed to the second round of the repechage contest. If the finalists for the lower branch did not compete in qualifications rounds, then there was no contestant 5, and contestant 6 was also byed to the second round of the repechage contest.

Note that in each bout of the repechage contests, the contestant wearing red had been eliminated "earlier" in the main tournament tree than the contestant wearing blue. If there is a link between the stage at which a contestant was eliminated and his skill, then this may potentially introduce a bias towards wins by blue. For example, in the outermost round of the repechage contest for the upper branch, the red-wearing contestant (1) had been eliminated in the qualifications round (round of 32), the blue-wearing contestant (2) in the eighth-final round (round of 16). The winner of this bout (3) wore red in the next bout; the blue-wearing contestant in this bout had reached the quarter-final round (round of 8). The winner of this bout wore red in the next bout; the blue-wearing contestant in this bout (4) had reached the semi-final round (round of 4). We note that there is no evidence of such a bias in the data (Greco-Roman wrestling: Section S3.2.3; free-style wrestling: Section S3.2.4), possibly due to the small number of rounds in each tournament that are potentially affected.

##### S2.4.4 Comparison across years

In wrestling there were substantial changes between the 2004 and 2008 competitions, affecting the overall tournament structure. Even these changes do not, in themselves, invalidate extension of Hill & Barton's [1] approach to data relating to the 2008 competition. In fact, because Hill & Barton [1] excluded bouts in the elimination pools from analysis of the 2004 data (Section S2.4.1), and elimination pools did not feature in the 2008 tournaments, the 2008 dataset more than trebles the number of bouts available for analysis in each Greco-Roman wrestling and free-style wrestling (Sections S3.1 and S3.2).

In light of these changes, we determined which bouts to exclude as walkovers from the 2008 wrestling data by comparing possible and realized bout outcomes for the two competitions, as follows. This ensures consistency with the exclusion criteria implemented by Hill & Barton [1] for the 2004 wrestling data.

For the 2004 competition, possible bout outcomes are:

`EF` Victory by forfeit, the loser is not classified
`EV` Disqualification from all competition for violation of the rules
`EX` 3 cautions or violation of the rules
`E2` Both wrestlers are disqualified for violation of the rules
`PA` Injury default
`PO` Victory by points, the loser without technical points
`PP` Victory by points, the loser with technical points
`SP` Technical superiority, 10 points difference, the loser with points
`ST` Technical superiority, 10 points difference, the loser without points
`TO` Victory by fall

All types except `EF`, `EX` occur in the Greco-Roman wrestling data analysed by Hill & Barton [1] (i.e., excluding bouts in the elimination pools). Similarly, all types except `EF`, `EX`, `E2`, `SP` occur in the free-style wrestling data analysed (i.e., also excluding bouts in the elimination pools). In both cases, the bouts coded as won by walkover by Hill & Barton [1] are of type `EV` or `PA`.

For the 2008 competition, possible bout outcomes are:

`E2` Both wrestlers have been disqualified due to infringement of the rules
`EX` 3 cautions '0' due to error against the rules
`PP` Decision by points, the loser with technical points
`ST` Great superiority, a difference of 6 points, the loser without points
`VB` Victory by injury
`VT` Victory by fall
`EV` Disqualification from the whole competition due to infringement of the rules
`PO` Decision by points, the loser without technical point
`SP` Victory by technical superiority with the loser scoring technical points
`VA` Victory by withdrawal
`VF` Victory by forfeit

The only types that occur in the data are `VT`, `ST`, `PP`, `PO` for Greco-Roman wrestling, and `VT`, `VA`, `ST`, `SP`, `PP`, `PO` for free-style wrestling. Of these, `VT`, `ST`, `SP`, `PP`, `PO` can be directly matched to types `TO`, `ST`, `SP`, `PP`, `PO` for the 2004 competition. Given that all of these occur in the 2004 data analysed by Hill & Barton [1] and were retained for analysis, we retained them in the 2008 data. Type `VA` cannot be directly matched to any type in the 2004 data. This corresponds to a walkover and was therefore excluded from the 2008 data.

#### S2.5 Summary

Competitions in the four sports analysed by Hill & Barton [1] were arranged as a single-elimination tournament for each weight class. Generally, in a single-elimination tournament contestants compete in pairs, with the winner of a bout proceeding to the next round in the competition tree. Above we have outlined features of the tournament structure and related aspects of relevance to the subsequent analyses; an overview is in Table S1. We have seen that details varied substantially — both across sports and, with the exception of boxing, within sports between the 2004 and 2008 competitions. We have also outlined the implications of this variation for extension of Hill & Barton's [1] approach from the 2004 to the 2008 data.

Table S1: Overview of relevant features of the tournament structure and related aspects in boxing (BOX), taekwondo (TKD), Greco-Roman wrestling (GRW), and free-style wrestling (FSW), at the 2004 Athens and 2008 Beijing Olympics. Included are the size of the outermost round of the main tournament trees, the average completeness fraction $\rho$ across weight classes, the relative placement of byes in the main tournament trees, and the color assigned to the contestant at the top of the bracket in each bout.

| Year | Sport | Outermost round | $\rho$ | Stacking of byes | Color |
|---|---|---|---|---|---|
| 2004 | BOX | 16 or 32 | 0.79 | Upper sub-tree | Red |
| 2004 | TKD | 16 | 1.00 | N/A | Blue |
| 2004 | GRW | 8 | 0.61 | Lower sub-tree | Red |
| 2004 | FSW | 8 | 0.68 | Lower sub-tree | Red |
| 2008 | BOX | 16 or 32 | 0.79 | Upper sub-tree | Red |
| 2008 | TKD | 16 | 1.00 | N/A | Blue |
| 2008 | GRW | 32 | 0.24 | Upper sub-tree | Red |
| 2008 | FSW | 32 | 0.24 | Upper sub-tree | Red |

Overall, through careful matching of the 2004 and 2008 data, we conclude that the two datasets are fully equivalent for the purpose of testing Hill & Barton's [1] hypothesis. Crucially, in all sports in both years, initial seeding of the main competition tree was based on a random draw. Seeding based on skill — a factor proposed as a potential source of bias towards wins by one color over the other in single-elimination tournaments [2, 3] — does not apply to these competitions, and it can therefore be ruled out as the explanation for any bias observed in the data.

Seeding based on skill is used to ensure even strength throughout the competition draw, for example to avoid two top-ranked contestants meeting in an early round, resulting in one of them being eliminated prematurely. Its introduction in boxing (Section S2.2.1) and in taekwondo (Section S2.3.1) at the 2012 London Olympics prevents extension of Hill & Barton's [1] approach to this competition.

### S3 Data for the sports analysed

#### S3.1 2004 Athens Olympics

The data were obtained from the supplementary information of Hill & Barton [1] (file `435293a-s1.xls`) and exported to `csv` format.

##### S3.1.1 Boxing

The data include all rounds. We excluded entries marked `Walk Over` in the column `Method of Win` (five total; three resulting in a win by red, two resulting in a win by blue).

The total number of entries for analysis is $n_{\mathrm{tot}} = 267$ bouts, with $n_{\mathrm{red}} = 147$ resulting in a win by red, $n_{\mathrm{blue}} = 120$ resulting in a win by blue.

##### S3.1.2 Taekwondo

The data include all rounds. We excluded entries marked `Withdrawn` in the column `Method of Win` (five total; two resulting in a win by red, three resulting in a win by blue).

The total number of entries for analysis is $n_{\mathrm{tot}} = 75$ bouts, with $n_{\mathrm{red}} = 43$ resulting in a win by red, $n_{\mathrm{blue}} = 32$ resulting in a win by blue.

##### S3.1.3 Greco-Roman wrestling

The data include only the qualification round, semi-finals, 1–2 final, 3–4 final, and 5–6 final (i.e., bouts from the elimination pools were not included; Section S2.4.1). We excluded entries marked `Yes` in the column `Won by Walkover` (three total; zero resulting in a win by red, three resulting in a win by blue).

The total number of entries for analysis is $n_{\mathrm{tot}} = 48$ bouts, with $n_{\mathrm{red}} = 25$ resulting in a win by red, $n_{\mathrm{blue}} = 23$ resulting in a win by blue.

##### S3.1.4 Free-style wrestling

The data include only the qualification round, semi-finals, 1–2 final, 3–4 final, and 5–6 final (i.e., bouts from the elimination pools were not included; Section S2.4.1). We excluded entries marked `Yes` in the column `Won by Walkover` (three total; one resulting in a win by red, two resulting in a win by blue).

The total number of entries for analysis is $n_{\mathrm{tot}} = 51$ bouts, with $n_{\mathrm{red}} = 27$ resulting in a win by red, $n_{\mathrm{blue}} = 24$ resulting in a win by blue.

##### S3.2 2008 Beijing Olympics

Results books were obtained from the LA84 Foundation archive at `http://library.la84.org/6oic/OfficialReports/2008/` (accessed on 2013-08-01; file `2008Results_Book1.pdf` for boxing, file `2008Results_Book2.pdf` for taekwondo, wrestling). The relevant data were entered manually and exported to `csv` format.

###### S3.2.1 Boxing

The data include all rounds. We excluded entries marked `WO` (walkover) in the column `method` (two total; zero resulting in a win by red, two resulting in a win by blue).

The total number of entries for analysis is $n_{\text{tot}} = 270$ bouts, with $n_{\text{red}} = 133$ resulting in a win by red, $n_{\text{blue}} = 137$ resulting in a win by blue.

###### S3.2.2 Taekwondo

The data include all rounds. We excluded entries marked `WDR` (withdrawn) in the column `method` (one total; zero resulting in a win by red, one resulting in a win by blue).

The total number of entries for analysis is $n_{\text{tot}} = 75$ bouts, with $n_{\text{red}} = 38$ resulting in a win by red, $n_{\text{blue}} = 37$ resulting in a win by blue.

###### S3.2.3 Greco-Roman wrestling

The data include all rounds. No entries were excluded, because none corresponding to walkovers are represented in the data (Section S2.4.4).

The total number of entries for analysis is $n_{\text{tot}} = 164$ bouts, with $n_{\text{red}} = 80$ resulting in a win by red, $n_{\text{blue}} = 84$ resulting in a win by blue.

Of the 32 entries corresponding to bouts in repechage and bronze rounds, 17 ended in a win by red, 15 in a win by blue (one-sided binomial test, $H_0 : f_{\text{blue}} \leq 0.5; H_{\text{A}} : f_{\text{blue}} > 0.5$, $p = 0.702$). There is thus no evidence of a bias towards wins by blue in the repechage contests (Section S2.4.3).

###### S3.2.4 Free-style wrestling

The data include all rounds. We excluded entries marked `VA` (victory by withdrawal; Section S2.4.4) in the column `method` (one total; one resulting in a win by red, zero resulting in a win by blue).

The total number of entries for analysis is $n_{\text{tot}} = 164$ bouts, with $n_{\text{red}} = 67$ resulting in a win by red, $n_{\text{blue}} = 97$ resulting in a win by blue.

Of the 33 entries corresponding to bouts in repechage and bronze rounds, 13 ended in a win by red, 20 in a win by blue (one-sided binomial test, $H_0 : f_{\text{blue}} \leq 0.5; H_{\text{A}} : f_{\text{blue}} > 0.5$, $p = 0.148$). There is thus no evidence of a bias towards wins by blue in the repechage contests (Section S2.4.3).

### S4 Replication

Hill & Barton [1] reported two sets of results for data relating to the 2004 Athens Olympics. The first set underpinned the main claim of an effect of red on the outcomes of Olympic combat sports. The analysis was in two parts. First, in each of the four sports, over 50% of the bouts resulted in a win by red, with a fraction statistically significantly different from 0.5 in the data aggregated over the four sports ($\chi^2 = 4.19$, d.f. = 1, $p = 0.041$). Second, further aggregation of the data across the four sports by round or by weight class revealed a consistent pattern: in both cases, the fraction including over 50% of wins by red was statistically significantly different from 0.5. Specifically, the four sports comprised a total of 21 rounds: of these, 16 presented a majority of wins by red, and only four a majority of wins by blue (sign test, $p = 0.012$). Similarly, of the 29 weight classes across the four sports, 19 presented a majority of wins by red, and only six a majority of wins by blue (sign test, $p = 0.015$).

The second set of results underpinned a corollary of the main claim — namely, that "the red advantage will determine the outcome only in relatively symmetric contests" [1, p. 293]. To evaluate this prediction, a final analysis divided the data aggregated over the four sports into four groups, based on the difference in the number of points for the two contestants at the end of each bout. The points difference was taken to indicate the degree of asymmetry in competitive ability. In the three most symmetric groups, over 50% of the bouts were won by red, with a fraction statistically significantly different from 0.5 in the most symmetric of the three (no asymmetries: $\chi^2 = 6.07$, d.f. = 1, $p = 0.014$; small asymmetries: $\chi^2 = 2.21$, d.f. = 1, $p = 0.14$; medium asymmetries: $\chi^2 = 0.47$, d.f. = 1, $p = 0.50$). In the least symmetric of the four groups, over 50% of the bouts were won by blue (large asymmetries: $\chi^2 = 0.21$, d.f. = 1, $p = 0.64$).

Attentive readers may have identified a number of issues with the analyses, as well as ample potential to employ "researcher degrees of freedom" [4] to obtain "evidence", in the form of statistically significant results, consistent with the hypothesis. This raises the possibility that the analyses were data-dependent — namely, that a different set of tests would have been performed on a different dataset [5]. In light of this, and to pre-empt concerns that any discrepancies between our results and those reported by Hill & Barton [1] hinge on specific analytic decisions, we proceed as follows. We begin by re-deriving the analytical approach implemented by Hill & Barton [1], with the two-fold aim to (i) reproduce the results for the 2004 data, and (ii) conduct an exact replication using the 2008 data (*sensu* Peng [6]; Section S4.1). In the process, we uncover several shortcomings with the approach, affecting the results underpinning both the main claim (Section S4.1.1) and its corollary (Section S4.1.2). Next, we devise the simplest alternative analytical approach to apply to both datasets — mirroring Hill & Barton's [1] as closely as possible, while addressing several of its key shortcomings (Section S4.2). The aim here is to show that they can be readily addressed. Therefore, the analytic decisions leading to them cannot be justified on the grounds of simplicity.

#### S4.1 Original analytical approach

##### S4.1.1 Main claim

**Reproduction (2004 Athens Olympics)** Table S2 reproduces the first part of the first set of results reported by Hill & Barton [1], following the analytical approach they applied to the 2004 data. The fraction of bouts won by red, $f_{\text{red}} = n_{\text{red}}/n_{\text{tot}}$, is greater than 50% in each of the four sports, but the only statistically significant result at the $\alpha = 0.05$ level is for the data aggregated over the four sports. Even this result is not robust, however: just one additional win by blue would tip the $p$-value over the significance threshold.

Table S2: Number of bouts won by red ($n_{\text{red}}$), total number of bouts ($n_{\text{tot}}$), fraction of bouts won by red ($f_{\text{red}} = n_{\text{red}}/n_{\text{tot}}$), and $\chi^2$ test results ($H_0 : f_{\text{red}} = 0.5; H_A : f_{\text{red}} \neq 0.5$; $\alpha = 0.05$) in data for boxing (BOX), taekwondo (TKD), Greco-Roman wrestling (GRW), free-style wrestling (FSW), and aggregated (ALL), at the 2004 Athens Olympics.

| Sport(s) | $n_{\text{red}}$ | $n_{\text{tot}}$ | $f_{\text{red}}$ | $\chi^2$ | d.f. | $p$ |
|---|---|---|---|---|---|---|
| BOX | 147 | 267 | 0.551 | 2.73 | 1 | 0.098 |
| TKD | 43 | 75 | 0.573 | 1.61 | 1 | 0.204 |
| GRW | 25 | 48 | 0.521 | 0.08 | 1 | 0.773 |
| FSW | 27 | 51 | 0.529 | 0.18 | 1 | 0.674 |
| ALL | 242 | 441 | 0.549 | 4.19 | 1 | 0.041 |

In any case, as the $\chi^2$ goodness-of-fit tests are performed in parallel on the same dataset, the analysis involves multiple hypothesis testing, with potentially serious consequences for the purpose of evaluating the set of results as a whole [7] (Section S4.2). Furthermore, we note that the $\chi^2$ test rests on an asymptotic approximation that only holds for large sample sizes. The binomial test is exact and correct for all sample sizes, and in its one-sided form it provides a more accurate representation of Hill & Barton's [1] prediction (i.e., $H_A : f_{\text{red}} > 0.5$ vs. $H_A : f_{\text{red}} \neq 0.5$). A final issue is that the data present dependencies linked to the tournament structure for the sports analysed, in violation of the independence assumption of standard hypothesis testing (Section S4.3).

Table S3 reproduces the second part of the first set of results reported by Hill & Barton [1], again following the analytical approach they applied to the 2004 data. The sign test used here is equivalent to a two-sided binomial test ($H_A : f_{\text{red}} \neq 0.5$), where the number of successes is the number of rounds (or weight classes) with a majority of wins by red, $n_{\text{red}}$, and the number of trials is the number of rounds (or weight classes) with a majority of wins by red or by blue, $n_{\text{red}} + n_{\text{blue}} < n_{\text{tot}}$. That is, rounds (or weight classes) with an equal number of wins by red and by blue are excluded from analysis. For both rounds and weight classes, the fraction presenting a majority of wins by red, $f_{\text{red}} = n_{\text{red}}/(n_{\text{red}} + n_{\text{blue}})$, is statistically significantly different from 0.5 at the $\alpha = 0.05$ level. However, neither result is robust: switching just two of the rounds (or weight classes) with a majority of wins by red to a majority of wins by blue would tip the $p$-value over the significance threshold.

Table S3: Number of rounds and weight classes with a majority of wins by red ($n_{\text{red}}$) or by blue ($n_{\text{blue}}$), total number rounds and weight classess ($n_{\text{tot}}$), and sign test results ($H_0 : f_{\text{red}} = 0.5; H_{\text{A}} : f_{\text{red}} \neq 0.5$; $\alpha = 0.05$) in data aggregated over boxing, taekwondo, Greco-Roman wrestling, and free-style wrestling, at the 2004 Athens Olympics. Note that $f_{\text{red}} = n_{\text{red}}/(n_{\text{red}} + n_{\text{blue}})$, with $n_{\text{red}} + n_{\text{blue}} < n_{\text{tot}}$.

| Test | $n_{\text{red}}$ | $n_{\text{blue}}$ | $n_{\text{tot}}$ | $p$ |
|---|---|---|---|---|
| Rounds | 16 | 4 | 21 | 0.012 |
| Weight classes | 19 | 6 | 29 | 0.015 |

The fact that the results are not robust is noteworthy, given specific analytic decisions implemented by Hill & Barton [1]. For example, rounds and weight classes can vary greatly in number of bouts, both within and between sports (Section S2). Consequently, the probability that different rounds or weight classes end with a majority of wins by one color, or with no majority, is not uniform. It is therefore not clear whether it is meaningful to aggregate them across sports. Furthermore, there is arbitrariness in the coding of rounds for analysis. For instance, Hill & Barton [1] collapsed all bouts in the taekwondo repechage contests (Section S2.3.3) into one round ("repechage"), except the final bouts of the contests, which they treated as a separate round ("bronze"). Finally, rounds and weight classes with an equal number of wins by red and by blue provide relevant information for the purpose of evaluating the prediction, yet Hill & Barton [1] excluded them from analysis. Failure to justify these decisions, coupled with results that are not robust, raises concerns that the analysis was, implicitly or explicitly, data-dependent [5] — effectively exploiting researcher degrees of freedom to obtain "findings" consistent with the hypothesis [4]. The probability of committing a Type I error (rejecting the null hypothesis, when the null hypothesis is in fact true — i.e., a false positive) is thus higher than the specified rate $\alpha = 0.05$ [4, 5].

We also note that, as above, a one-sided test provides a more accurate representation of Hill & Barton's [1] prediction (i.e., $H_{\text{A}} : f_{\text{red}} > 0.5$ vs. $H_{\text{A}} : f_{\text{red}} \neq 0.5$), and that we are in a setting of multiple hypothesis testing (Section S4.2), with dependencies in the data linked to the tournament structure for the sports analysed (Section S4.3).

**Replication (2008 Beijing Olympics)** Tables S4 and S5 summarize the $\chi^2$ and sign test results for the 2008 data, reported here as an exact replication of Hill & Barton's [1] analysis (Section S2.5). Thus, the shortcomings with the analytical approach outlined above extend to analysis of this independent dataset.

Qualitatively, the pattern observed in these data is reversed compared to the 2004 data. The fraction of bouts won by blue is greater than 50% in all the sports except taekwondo, which presents an "excess" of only one win by red (Table S4). Consistently, more rounds and weight classes present a majority of wins by blue than a majority of wins by red (Table S5).

Table S4: Number of bouts won by red ($n_{\text{red}}$), total number of bouts ($n_{\text{tot}}$), fraction of bouts won by red ($f_{\text{red}} = n_{\text{red}}/n_{\text{tot}}$), and $\chi^2$ test results ($H_0 : f_{\text{red}} = 0.5; H_{\text{A}} : f_{\text{red}} \neq 0.5$; $\alpha = 0.05$) in data for boxing (BOX), taekwondo (TKD), Greco-Roman wrestling (GRW), free-style wrestling (FSW), and aggregated (ALL), at the 2008 Beijing Olympics.

| Sport(s) | $n_{\text{red}}$ | $n_{\text{tot}}$ | $f_{\text{red}}$ | $\chi^2$ | d.f. | $p$ |
|---|---|---|---|---|---|---|
| BOX | 133 | 270 | 0.493 | 0.06 | 1 | 0.808 |
| TKD | 38 | 75 | 0.507 | 0.01 | 1 | 0.908 |
| GRW | 80 | 164 | 0.488 | 0.10 | 1 | 0.755 |
| FSW | 67 | 164 | 0.409 | 5.49 | 1 | 0.019 |
| ALL | 318 | 673 | 0.473 | 2.03 | 1 | 0.154 |

Table S5: Number of rounds and weight classes with a majority of wins by red ($n_{\text{red}}$) or by blue ($n_{\text{blue}}$), total number rounds and weight classess ($n_{\text{tot}}$), and sign test results ($H_0 : f_{\text{red}} = 0.5; H_{\text{A}} : f_{\text{red}} \neq 0.5$; $\alpha = 0.05$) in data aggregated over boxing, taekwondo, Greco-Roman wrestling, and free-style wrestling, at the 2008 Beijing Olympics. Note that $f_{\text{red}} = n_{\text{red}}/(n_{\text{red}} + n_{\text{blue}})$, with $n_{\text{red}} + n_{\text{blue}} < n_{\text{tot}}$.

| Test | $n_{\text{red}}$ | $n_{\text{blue}}$ | $n_{\text{tot}}$ | $p$ |
|---|---|---|---|---|
| Rounds | 8 | 13 | 25 | 0.383 |
| Weight classes | 11 | 17 | 29 | 0.345 |

Across the seven hypothesis tests, the only statistically significant result at the $\alpha = 0.05$ level is for bouts in free-style wrestling, but the deviation from the null of $f_{\text{red}} = 0.5$ is in the opposite direction than predicted (Table S4). It is obvious in this case why, as discussed above, a one-sided test (i.e., $H_{\text{A}} : f_{\text{red}} > 0.5$ vs. $H_{\text{A}} : f_{\text{red}} \neq 0.5$) is more appropriate.

###### S4.1.2 Corollary

Due to ambiguities in processing of the data, we were able to reproduce the second set of results reported by Hill & Barton [1] only approximately (not shown). As outlined above, this set underpinned a corollary of the main claim — namely, that "the red advantage will determine the outcome only in relatively symmetric contests" [1, p. 293]. The 2004 data were thus aggregated over the four sports and divided into four groups, based on the difference in the number of points for the two contestants in each bout, corresponding to different degrees of asymmetry in competitive ability. Our attempt to reproduce the results failed in re-deriving the exact procedure Hill & Barton [1] used to obtain the groups, following the description they provided in the supplementary information (file `435293a-s2.doc`).

In any case, based on close examination of the results as reported, we contend that they must be discounted, for two reasons. First, the underlying analysis is compromised by a series of shortcomings analogous to the ones discussed in Section S4.1.1. Second, any attempt to extract information about the degree of asymmetry in competitive ability from the points difference is critically flawed. We elaborate on both issues below.

Hill & Barton's [1] analysis involved dividing the data aggregated over the four sports into four groups, with a $\chi^2$ goodness-of-fit test performed on each. Thus, again we are in a setting of multiple hypothesis testing (Section S4.2). Given the relatively small size of the subsets (on the order of 110 bouts each), and the specific prediction (i.e., $H_\mathrm{A} : f_\mathrm{red} > 0.5$ vs. $H_\mathrm{A} : f_\mathrm{red} \neq 0.5$), a one-sided binomial test is more appropriate than the $\chi^2$ test, for reasons outlined in Section S4.1.1. Furthermore, the subsets retain the dependencies of the full dataset (Section S4.3). Another issue is with Hill & Barton's [1] interpretation of the results, which rested on comparison of the levels of statistical significance across the groups. It is well established that comparisons of this sort can be misleading [8].

Ignoring concerns about researcher degrees of freedom, these shortcomings could be readily addressed by implementing an alternative analytical approach. However, any such effort is curbed by a crucial issue with the data. Specifically, using the difference in points scored by the two contestants in a bout as a proxy for asymmetry in competitive ability, and aggregating the information over the four sports, is an uncontrolled procedure with unknown properties. The results yielded by such a procedure are thus of difficult interpretation, at best; at worst, they are meaningless. This is because point-scoring systems vary greatly across the four sports analysed, and the number of points scored is not "linear" with respect to skill. For example, boxing presents the simplest scoring system across the four sports. Broadly, one point is assigned for a punch meeting specific requirements that lands on the opponent's head or torso. However, judges rely on additional considerations (e.g., better style, better defense) to break ties. Thus, the assumption of a "linear" relationship between the number of points scored by a contestant and his skill is questionable even in this case.

The assumption is even less tenable for the scoring systems of taekwondo and wrestling, which assign a different number of points for different "actions". In taekwondo, an additional complication is the deduction of penalty points, which may result in a contestant ending a bout with an overall negative score. Even when the overall scores are positive for both contestants in a bout, interpretation of the points difference as a proxy for asymmetry in competitive ability is problematic, invalidating comparison with the other sports. In wrestling, a further difficulty arises from the distinction between technical vs. classification points. Technical points are the actual points awarded to the contestants during a bout; classification points are awarded at the end of the bout based on a number of considerations — the number of technical points scored in the bout being one of them! Hence, it is not clear that summing technical and classification points, as Hill & Barton [1] did, is in any way meaningful — also invalidating comparison with the other sports.

#### S4.2 Alternative analytical approach

Here we implement an alternative approach to replicate the first set of results reported by Hill & Barton [1], which underpinned the main claim of an effect of red on the outcomes of Olympic combat sports (Section S4.1.1). As noted above, we devise the simplest analytical approach to apply to both datasets — mirroring Hill & Barton's [1] approach as closely as possible, but addressing several of its key shortcomings. The aim is to show that these are readily addressed within an analogous paradigm. Therefore, the analytical decisions leading to them cannot be justified, for example on the grounds of simplicity.

We emphasize that the alternative approach does not address the issue of dependencies in the data linked to the tournament structure for the sports analysed — this issue is more subtle, requiring separate treatment (Section S4.3). We describe the approach in detail here, because it will be applied again in a final analysis of the data which will address this oustanding issue (Section S6.3).

We use a series of one-sided binomial tests ($H_0 : f_{\text{red}} \leq 0.5; H_{\text{A}} : f_{\text{red}} > 0.5$; $\alpha = 0.05$), applied to (i) the fraction of bouts won by red, separately for individual sports and aggregated over the four sports by year, and (ii) the fraction of rounds and weight classes with a majority of wins by red, aggregated over the four sports by year. Note that $f_{\text{red}} = n_{\text{red}}/n_{\text{tot}}$ in all cases — that is, unlike in Hill & Barton's [1] analysis, rounds and weight classes with an equal number of wins by red and by blue are included, as they provide relevant information for the purpose of evaluating the prediction (Section S4.1.1).

The results are in Table S6. In addition to the raw $p$-values, the table includes the $100(1-2\alpha)\% = 90\%$ confidence intervals for $f_{\text{red}}$, calculated using Wilson's [9] method [10], and the $p$-values adjusted for multiple hypothesis testing using four methods (Bonferroni [11, 12], Holm [13], Benjamini & Hochberg [14], and Benjamini & Yekutieli [15]). Crucially, there is substantial variation in the values of $f_{\text{red}}$, both within and across years. Focusing on the raw $p$-values, only two of the 14 tests return results that are statistically significant at the $\alpha = 0.05$ level, both relating to the 2004 data. The variables involved are (i) the fraction of bouts won by red aggregated over the four sports, and (ii) the fraction of rounds with a majority of wins by red, aggregated over the four sports.

The result relating to the second variable is of difficult interpretation, because the number of bouts varies across rounds, both within and between sports (Section S4.1.1). Additionally, the two variables are not independent, in the sense that as one increases, so does the other. A related issue is that we are in a setting of multiple hypothesis testing, with potentially serious consequences for the purpose of evaluating the set of results as a whole [7]. Specifically, we may be committing a Type I error (rejecting the null hypothesis, when the null hypothesis is in fact true — i.e., a false positive), at a rate higher than the specified Type I error probability $\alpha = 0.05$. We have chosen this value as the acceptable maximum probability of a Type I error for each test to mirror Hill & Barton's [1] analytical approach (Section S4.1.1). But in a setting of multiple hypothesis testing, the probability of committing at least some Type I errors grows with the number of tests [7].

There is a rich literature on corrections for multiple hypothesis testing, but no universally applicable, or accepted, method — this is an active area of research. Two established approaches are to control the family-wise error rate, and to control the false discovery rate [7]. For completeness, we report the $p$-values adjusted for multiple hypotheses testing using both approaches (Table S6).

The methods of Bonferroni [11, 12] and Holm [13] control the family-wise error rate. This approach attempts to limit the probability of even a single Type I error within the family of tests. The methods of Benjamini & Hochberg [14] and Benjamini & Yekutieli [15] control the false discovery rate within the set of tests. A false discovery corresponds to incorrectly rejecting the null hypothesis, and the false discovery rate is the expected proportion of incorrect rejections out of all rejections. This approach focuses on limiting the expected proportion of false discoveries, out of all discoveries, within the set of tests.

The family-wise error rate is a more stringent condition than the false discovery rate, corresponding to lower "tolerance" of Type I errors. This comes at the cost of statistical power, possibly resulting in Type II errors (not rejecting the null hypothesis, when the alternative hypothesis is true instead — i.e., a false negative). Generally, then, controlling the family-wise error rate is a conservative procedure — more so than controlling the false discovery rate [14, 15]. Bonferroni [11, 12] is the most conservative of the methods we apply. It is has lower statistical power than Holm [13], which in turn has lower power than the two false discovery rate procedures. Of these, Benjamini & Hochberg [14] is less conservative, but it makes specific assumptions about the joint distribution of $p$-values, which are not required by Benjamini & Yekutieli [15]. Therefore, we apply both.

The adjusted $p$-values have different interpretations for the two approaches. For methods that control the family-wise error rate, the adjusted $p$-value for a test is the smallest family-wise level of $\alpha$ at which the corresponding null hypothesis would be rejected [16]. For methods that control the false discovery rate, it is the lowest level of the false discovery rate at which the hypothesis would be first included in the set of rejected hypotheses [17].

For the 14 tests in Table S6, we can begin by examining the results of the Bonferroni [11, 12] and Holm [13] methods. Controlling the family-wise error rate at level $\alpha = 0.05$ — that is, evaluating the adjusted $p$-values against this threshold — produces no statistically significant results for either method. Closer inspection of the adjusted $p$-values shows that a family-wise error rate $\alpha \geq 0.32$ (under Bonferroni [11, 12]) or $\alpha \geq 0.29$ (under Holm [13]) would be required to reject the two null hypotheses rejected with no adjustment. These values correspond to a high probability of committing a Type I error.

Table S6: Results of one-sided binomial tests in data for boxing (BOX), taekwondo (TKD), Greco-Roman wrestling (GRW), free-style wrestling (FSW), and aggregated over the four sports (ALL), at the 2004 Athens and 2008 Beijing Olympics. Tests denoted “bouts” compare the number of bouts won by red, $n_{\mathrm{red}}$, to the total number of bouts, $n_{\mathrm{tot}}$. Tests denoted “rounds” and “weight classes” compare the number with a majority of wins by red, $n_{\mathrm{red}}$, to the total number, $n_{\mathrm{tot}}$. In all cases, $f_{\mathrm{red}} = n_{\mathrm{red}}/n_{\mathrm{tot}}$. Included are the raw $p$-values ($H_0 : f_{\mathrm{red}} \leq 0.5; H_{\mathrm{A}} : f_{\mathrm{red}} > 0.5; \alpha = 0.05$), the 90% confidence intervals for $f_{\mathrm{red}}$, and the $p$-values adjusted for multiple hypothesis testing using four methods (B: Bonferroni [11, 12]; H: Holm [13]; BH: Benjamini & Hochberg [14]; BY: Benjamini & Yekutieli [15]).

| Year | Test | Sport(s) | $n_{\mathrm{red}}$ | $n_{\mathrm{tot}}$ | $f_{\mathrm{red}}$ | 90% CI | $p$ | | | | |
|---|---|---|---|---|---|---|---|---|---|---|---|
| | | | | | | | Raw | B | H | BH | BY |
| 2004 | Bouts | BOX | 147 | 267 | 0.551 | 0.500–0.600 | 0.056 | 0.780 | 0.668 | 0.238 | 0.774 |
| 2004 | Bouts | TKD | 43 | 75 | 0.573 | 0.478–0.663 | 0.124 | 1.000 | 1.000 | 0.347 | 1.000 |
| 2004 | Bouts | GRW | 25 | 48 | 0.521 | 0.404–0.635 | 0.443 | 1.000 | 1.000 | 0.875 | 1.000 |
| 2004 | Bouts | FSW | 27 | 51 | 0.529 | 0.416–0.640 | 0.390 | 1.000 | 1.000 | 0.875 | 1.000 |
| 2004 | Bouts | ALL | 242 | 441 | 0.549 | 0.510–0.587 | 0.023 | 0.318 | 0.295 | 0.159 | 0.516 |
| 2004 | Rounds | ALL | 16 | 21 | 0.762 | 0.585–0.879 | 0.013 | 0.186 | 0.186 | 0.159 | 0.516 |
| 2004 | Weight classes | ALL | 19 | 29 | 0.655 | 0.502–0.781 | 0.068 | 0.952 | 0.748 | 0.238 | 0.774 |
| 2008 | Bouts | BOX | 133 | 270 | 0.493 | 0.443–0.542 | 0.620 | 1.000 | 1.000 | 0.913 | 1.000 |
| 2008 | Bouts | TKD | 38 | 75 | 0.507 | 0.413–0.600 | 0.500 | 1.000 | 1.000 | 0.875 | 1.000 |
| 2008 | Bouts | GRW | 80 | 164 | 0.488 | 0.424–0.552 | 0.652 | 1.000 | 1.000 | 0.913 | 1.000 |
| 2008 | Bouts | FSW | 67 | 164 | 0.409 | 0.347–0.473 | 0.992 | 1.000 | 1.000 | 0.992 | 1.000 |
| 2008 | Bouts | ALL | 318 | 673 | 0.473 | 0.441–0.504 | 0.929 | 1.000 | 1.000 | 0.992 | 1.000 |
| 2008 | Rounds | ALL | 8 | 25 | 0.320 | 0.191–0.484 | 0.978 | 1.000 | 1.000 | 0.992 | 1.000 |
| 2008 | Weight classes | ALL | 11 | 29 | 0.379 | 0.247–0.532 | 0.932 | 1.000 | 1.000 | 0.992 | 1.000 |

Turning to the results of the Benjamini & Hochberg [14] and Benjamini & Yekutieli [15] methods, controlling the false discovery rate at 5%, 10%, or 15% — that is, evaluating the adjusted $p$-values against these thresholds — produces no statistically significant results for either method. As before, closer inspection of the adjusted $p$-values provides additional information. Focusing on the two null hypotheses rejected with no adjustment, a false discovery rate of at least 16% (under Benjamini & Hochberg [14]) or 52% (under Benjamini & Yekutieli [15]) would be required for these to be included in the set of rejections. In other words, we can expect $2 \times 0.159 = 0.3$ (under Benjamini & Hochberg [14]) or $2 \times 0.516 = 1$ (under Benjamini & Yekutieli [15]) of the two rejections to be false discoveries.

#### S4.3 Dependencies in the data

Statistical procedures like the $\chi^2$, sign, and binomial tests, as applied in Sections S4.1 and S4.2, rest on the standard assumption that observations are independent and identically distributed. However, the bouts in a single-elimination tournament cannot be considered independent observations for the purpose of hypothesis testing: it is a defining feature of this type of tournament that winning contestants compete in multiple rounds (e.g., as they proceed to the final; Section S2). Specific features of the tournament structure may lead to associations between a contestant's skill, and hence the probability that he wins a given bout, and the color he wears. This can result in bias towards wins by one color over the other in the data aggregated over multiple rounds, competitions, and so on [2].

Determining the effect of any such bias from observational data alone is problematic. One approach in the literature has been to exclude from analysis sub-sets of bouts that are believed to be most affected by the potential bias [e.g., 2, 18, 19, for related investigation of the effect of blue on the outcomes of judo tournaments]. Doing so reduces statistical power, however, making it difficult to interpret the results as a whole. Furthermore, the bias may "percolate" across rounds in a tournament. Consequently, it is not clear that excluding specific sub-sets of bouts addresses the issue.

An alternative approach would be to incorporate into the analysis information about variance in skill among the contestants. For the four sports analysed here, the only source of available data involves the number of points for the two contestants in each bout. We have discussed in Section S4.1.2 the impediments to using these data.

In theory, it may be possible to include into the analysis contestant-specific parameters, to model the correlation structure induced by contestants featuring in multiple rounds [e.g., 2]. In practice, however, the tournament trees for the four sports analysed here are not sufficiently "deep" to provide enough information towards this (Section S2). Specifically, the deepest trees include only five rounds (i.e., the trees starting with a 32-contestant round). Consequently, the two contestants reaching the final on these trees compete in *at most* five bouts — less than five, if byes and/or walkovers are involved. All other contestants on these trees compete in four bouts or less. Trees of such depth are only found in some weight classes in boxing (Table S1). In the other weight classes in boxing, and in all weight classes in taekwondo and in wrestling, contestants compete in only three bouts or less on the main tournament tree. In taekwondo and in wrestling contestants may compete in additional bouts in the consolation rounds (i.e., the 3–4 and 5–6 final rounds in 2004 wrestling, and the repechage contests in taekwondo and in 2008 wrestling; Sections S2.3.3 and S2.4.3), but these rounds present several peculiar features compared to the main tournaments (Section S6.1.2). A related issue is that contestants only compete against each other once — that is, in a single bout throughout the tournament. This prevents inference of skill levels from the data through standard tools, such as the Bradley–Terry model of competition [20] (Section S5). It follows that simply adding data relating to other competitions, Olympic or otherwise [e.g., 2], is also not the solution — analogous issues would likely affect the additional data.

##### S4.4 Summary

Hill & Barton [1] predicted an effect of red on the outcomes of Olympic combat sports, which they tested against data for four sports (male divisions) at the 2004 Athens Olympics. They reported two sets of results. The first set focused on the fraction of bouts won by red, and on the fraction of rounds and weight classes with a majority of red wins. The second set focused on the fraction of bouts won by red across four groups, capturing different levels of asymmetry in the competitive ability of the contestants. The two sets of results underpinned, respectively, (i) the main claim of an effect of red, and (ii) its corollary that the effect only applies in relatively symmetric scenarios.

We began by reproducing (*sensu* Peng [6]) the first set of results (Section S4.1.1), which involved re-deriving the analytical approach Hill & Barton [1] applied to the 2004 data. Of the seven hypothesis tests, three returned statistically significant results, but significance hinged on only one or two observations in each case. Through this process, we identified a number of data-dependent analytic decisions that were not justified explicitly by Hill & Barton [1], indicating that a different set of tests may have been performed had different data been available [5]. Furthermore, we uncovered a number of shortcomings with the analytical approach, ranging from issues of test mis-specification to failure to adjust for multiple hypothesis testing and for dependencies in the data linked to the structure of the sports analysed. Overall, it seems likely that the work relied on so-called "researcher degrees of freedom" [4] — flexibility in processing and analysis of the data, and in interpretation and reporting of the results — to obtain "findings" consistent with the hypothesis.

Next, in Section S4.1.1 we performed an exact replication (*sensu* Peng [6]), by extending the analytical approach to equivalent data for the 2008 Beijing Olympics. We found that the first set of results reported by Hill & Barton [1] fails to replicate in this independent dataset. In fact, the qualitative pattern is reversed here compared to the 2004 data: in the 2008 data there is a preponderance of wins by blue.

In Section S4.1.2 we turned to the second set of results reported by Hill & Barton [1]. We found that analogous shortcomings and concerns apply in this case. The analysis involved four hypothesis tests, with the 2004 data aggregated over the four sports and divided into four groups, based on the difference in the number of points for the two contestants in each bout. Due to ambiguities in processing of the data — specifically, relating to the exact procedure Hill & Barton [1] used to obtain the groups — we were able to reproduce this set of results only approximately. In any case, from a close examination of the point-scoring systems for the four sports, we concluded that the premise underlying the analysis is critically flawed. That is, any attempt to extract information about variation in competitive ability from the points difference is compromised by subtleties of the scoring systems and their variation across the sports. It follows that the second set of results reported by Hill & Barton [1] must be discounted.

Finally, in Section S4.2 we replicated the first set of results reported by Hill & Barton [1] by implementing an alternative analytical approach. The aim was to devise the simplest approach to apply to both datasets — mirroring Hill & Barton's [1] approach as closely as possible, while addressing several of its shortcomings (Section S4.1.1).

The results show that there is substantial variation in the fraction of wins by red, both within and across years. For example, the fraction of bouts won by red ranges from 0.573 in 2004 taekwondo to 0.409 in 2008 free-style wrestling. Contrary to the prediction, then, the largest deviation from a 50% split observed in any sport in either year is towards wins by blue, not red. In itself, this variation invites caution in drawing inferences about the effect of color on the outcomes of Olympic combat sports. Consistently, even based on the raw $p$-values the null hypothesis of no effect of red is rejected in only two of 14 tests, both relating to the 2004 data. Inspection of the confidence intervals and of the $p$-values adjusted for multiple hypothesis testing suggests that these rejections are likely spurious.

A number of issues further complicate interpretation. For example, the two variables in these tests are not independent — effectively, they both capture the fraction of wins by red in 2004, such that as one variable increases, so does the other. One of the two variables is the fraction of rounds with a majority of wins by red, aggregated over the four sports. This variable is problematic in itself, due to variation in the number of bouts across rounds, both within and between sports (Section S4.1.1).

Finally, there is the possibility that the pattern observed in the data — with a predominance of wins by red in 2004 and blue in 2008 — is not simply the product of random variation. Rather, it may also reflect bias towards wins by one color over the other, arising from dependencies in the data linked to specific features of the tournament structure. As discussed in Section S4.1.1, the effect of any such bias is not determined using existing approaches in the literature, nor standard statistical procedures.

### S5 Mechanism

We discussed in Section S4 that dependencies in the data linked to the tournament structure for the sports analysed may result in bias towards wins by one color over the other. In this section we investigate the underlying mechanism to gain insight into the effect of the bias.

We begin by outlining the intuition behind the mechanism (Section S5.1), followed by mathematical analysis of a toy model (Section S5.2) and by numerical analysis of more general versions of the model (Section S5.3).

#### S5.1 Intuition

Recall from Section S2 that, in each bout, the contestant at the top of the bracket wore red in boxing and wrestling, blue in taekwondo (Table S1). Thus, the color worn by a contestant was linked to his relative position in the bout (top vs. bottom of the bracket), with the assignment procedure fixed throughout the tournament.

As we show below, bias towards wins by one color over the other arises from this assignment procedure, coupled with variance in contestant skill and with incompletness in the tournament tree. The underlying mechanism is asymmetric selection on contestant skill. Selection is an intrinsic feature of the tournament: effectively, the aim is to progressively eliminate contestants with relatively lower skill, retaining those with relatively higher skill as they proceed through the rounds. In an ideal scenario, the two contestants with the highest skill to enter the competition will face off in the final bout.[1] As the tournament progresses, then, this selective process shifts the distribution of contestant skill towards higher values. If the tournament tree is complete, the shift is symmetric in the upper and lower sub-trees. If the tournament tree is incomplete, the shift may be asymmetric.

Recall from Section S2 that two sources of incompleteness in a tournament tree are byes and walkovers. Both result in missing bouts, but whereas walkovers can occur anywhere on the tree, byes are always placed in the outermost round. In particular, byes can be drawn from the highest position in the round going down, or from the lowest position going up. These procedures result in byes stacked in the upper sub-tree or in the lower one, respectively. In one case there are more byes (and thus fewer contestants) in the upper sub-tree; in the other, there are more byes (and thus fewer contestants) in the lower sub-tree (Section S2.1).

As we show below, this patterning in the placement of byes results in weaker selection on contestant skill in the sub-tree with more byes (and thus fewer contestants). In other words, byes in the outermost round of the tree induce an asymmetry in the selective effect of bouts leading up to the final. As a result, the expected skill level of one contestant in the final will exceed his opponent's, making that contestant more likely to win. If the color

[1] In real tournaments, this pair may face off in an earlier round, with one being eliminated prematurely — hence the implementation of procedures such as seeding by skill (Section S2.5).

assignment procedure is as above, then the winner of the final will be more likely to wear a particular color.

This effect holds throughout the tournament. If the two sub-trees feeding into a bout include different numbers of contestants, the expected skill level of the contestant in one bout position (top vs. bottom of the bracket) will exceed his opponent's, making the contestant in that position more likely to win. Overall, then, there will be more wins by contestants wearing the color associated with that bout position, leading to *systematic bias* towards wins by that color over the other. This bias is systematic because, as we show below, its direction and magnitude can be predicted based on a few key factors — namely, the distribution of contestant skill, the color assignment procedure, the placement of the byes, and their number relative to the number of contestants entering the tournament.

Analogous reasoning applies to walkovers, with the crucial difference that, in this case, there is no pattern in the placement of the missing bouts with respect to each other and, ultimately, with respect to color. Consequently, whether walkovers do result in asymmetric selection on contestant skill, and whether this, in turn, results in bias towards wins by one color over the other, depends on the specific configuration. Thus, walkovers may induce *idiosyncractic bias* towards wins by one color over the other. This bias is idiosyncratic because its direction and magnitude are not easily predicted.

If a tree presents both systematic and idiosyncractic biases, due to byes and walkovers respectively, these may interact synergistically or antagonistically — their combined effect again depends on the specific configuration, such that direction and magnitude of the overall bias are not easily predicted. For simplicitly, then, the focus here is on the effect of byes, to the exclusion of both walkovers and bouts in the consolation rounds. More realistic scenarios, featuring all potential sources of bias (i.e., missing bouts due to byes and/or walkovers in the main tournament tree and in any consolation rounds), are investigated numerically in Section S6.2.

#### S5.2 Mathematical analysis

We first demonstrate the existence of the mechanism in a general setting (Section S5.2.1), and then we derive exact upper bounds on systematic bias towards wins by one color over the other in small tournaments (Section S5.2.2).

##### S5.2.1 General setting

The color assignment procedure translates into a fixed rule determining which color a contestant wears in a particular bout $X$, based on whether he entered $X$ from the top or bottom sub-tree rooted at $X$. Here a sub-tree is defined to include a particular bout $X$, the root of that sub-tree, and all the preceding bouts whose outcomes determined which contestants face each other at $X$. The number of contestants in the sub-tree is $n_X$, and the sub-tree is complete when $n_X$ is a power of 2. In this case, the length of a path from an

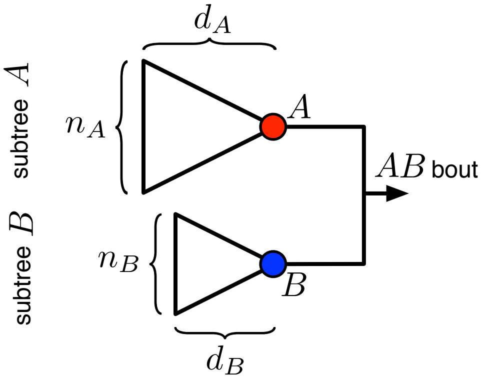

Figure S1: Diagram of a single-elimination tournament including $n = n_A + n_B$ contestants, with $n_A > n_B$. The winners $A$ and $B$ of the two sub-trees face off in a bout $AB$, in which $A$ wears red and $B$ wears blue. To reach this bout, they each won exactly $d_A$ and $d_B$ bouts.

outermost position to the root is $d_X$, and the sub-tree has depth $d = \log_2 n_X$ (corresponding to the number of rounds; Section S2.1). This implies that a contestant who reaches the root of the sub-tree has previously won exactly $d$ bouts (Figure S1).

For the general setting, we analyze the case of two sub-trees, $A$ and $B$, each including $n_A$ and $n_B$ contestants (Figure S1). We call $A$ and $B$ the top and bottom sub-trees of the $AB$ bout, and we use the fixed rule that the winners of the top and bottom sub-trees feeding into a bout wear red and blue, respectively, in the bout. Every tournament with any number of byes in the outermost round contains an $AB$ pair of sub-trees, hence demonstrating the existence of the mechanism in this general setting covers less abstract settings as well.

To decide which of two contestants $i, j$ wins a bout, we use the Bradley–Terry rule [20], i.e., $\Pr(i \text{ wins over } j) = \pi_i/(\pi_i + \pi_j)$, where $\pi$ is a contestant's skill level. The skill level of any particular contestant is fixed and given by $\pi \in \{\epsilon, 1\}$ with equal probability, where $\epsilon \approx 0$. This choice gives the maximum variance in the skill distribution, which is useful to illustrate the magnitude of the bias that can arise through asymmetric selection on skill. Other skill distributions yield similar results, but they require more complicated analytic calculations and/or numerical simulations.

**Base case** We begin with the base case of $n_A = n_B = 1$, which represents two contestants facing each other with no history of wins or losses in the tournament. There are four configurations of skill for $A$ and $B$, i.e., $\{(\epsilon, \epsilon), (\epsilon, 1), (1, \epsilon), (1, 1)\}$, all occurring with equal probability. In the $(\epsilon, \epsilon)$ configuration the winner of the $AB$ bout will have skill $\pi = \epsilon$. In the other three configurations the winner will have skill $\pi = 1$, because $\pi = 1$ always beats $\pi = \epsilon$. The probability that the $AB$ bout is won by the contestant with lower vs. higher skill is thus $\frac{1}{4}$ vs. $\frac{3}{4}$. Hence, the tournament effectively enhances the probability that the

Table S7: Probabilities and bout outcomes for the possible configurations of skill $\pi$ for contestants $A$ and $B$ in Figure S1, in the base case of $n_A = n_B = 1$.

| $\pi_A$ | $\pi_B$ | $\Pr(\pi_A, \pi_B)$ | $\pi_{\text{winner}}$ | Pr(red wins) |
|---|---|---|---|---|
| $\epsilon$ | $\epsilon$ | $1/2 \times 1/2 = 1/4$ | $\epsilon$ | 1/2 |
| $\epsilon$ | 1 | $1/2 \times 1/2 = 1/4$ | 1 | 0 |
| 1 | $\epsilon$ | $1/2 \times 1/2 = 1/4$ | 1 | 1 |
| 1 | 1 | $1/2 \times 1/2 = 1/4$ | 1 | 1/2 |

winner of the bout is the contestant with higher skill (Section S5.1). Table S7 provides a complete accounting of the four skill configurations for this base case, their likelihoods, and the probability of red winning conditioned on a particular configuration.

The total probability that $A$ (wearing red) wins the $AB$ bout is then given by

$$\begin{aligned}
\Pr(\text{red wins}) &= \sum_{\{\pi_A, \pi_B\}} \Pr(\text{red wins}) \Pr(\pi_A, \pi_B) \\
&= \frac{1}{2} \Pr(\epsilon, \epsilon) + \Pr(1, \epsilon) + \frac{1}{2} \Pr(1, 1) \\
&= \frac{1}{2}\left(\frac{1}{4}\right) + \frac{1}{4} + \frac{1}{2}\left(\frac{1}{4}\right) \\
&= \frac{1}{2} \ .
\end{aligned}$$

In the base case, then, the outcome is symmetric with respect to color, and the tournament will tend to select as the winner the contestant with higher skill.

**General case** Extending the reasoning to the more general case of $n_A, n_B > 1$, the only scenario in which contestant $A$ in the $AB$ bout will have skill $\pi_A = \epsilon$ is if all $n_A$ contestants in the $A$ sub-tree have skill $\epsilon$. Each contestant $i$ has skill $\pi_i = \epsilon$ with probability $\frac{1}{2}$, so this occurs with probability $2^{-n_A}$. In all other scenarios contestant $A$ in the $AB$ bout will have skill $\pi_A = 1$, and this must then occur with probability $1 - 2^{-n_A}$. The same calculation holds for contestant $B$.

The effect of the bouts in the $A$ sub-tree is thus to select a contestant with higher skill to the $AB$ bout, by shifting the distribution of skill $\pi_A$ towards higher values, and likewise for bouts in the $B$ sub-tree. We show next that, when $n_A = n_B$, the selective effect is equal in the $A$ and $B$ sub-trees, and the outcome of the $AB$ bout is symmetric with respect to color. But when $n_A \neq n_B$, the selective effect is weaker in the sub-tree with fewer contestants. With red and blue associated with the top and bottom sub-trees, respectively, this leads to a bias towards wins by red when $n_A > n_B$, and towards wins by blue when $n_A < n_B$. The magnitude of the bias can be calculated analytically in the toy model under study,

Table S8: Probabilities and bout outcomes for the possible configurations of skill $\pi$ for contestants $A$ and $B$ in Figure S1, in the general case of $n_A, n_B > 1$.

| $\pi_A$ | $\pi_B$ | $\Pr(\pi_A, \pi_B)$ | $\pi_{\text{winner}}$ | Pr(red wins) |
|---|---|---|---|---|
| $\epsilon$ | $\epsilon$ | $2^{-n_A} \times 2^{-n_B} = 2^{-(n_A+n_B)}$ | $\epsilon$ | 1/2 |
| $\epsilon$ | 1 | $2^{-n_A} \times (1 - 2^{-n_B}) = 2^{-n_A} - 2^{-(n_A+n_B)}$ | 1 | 0 |
| 1 | $\epsilon$ | $(1 - 2^{-n_A}) \times 2^{-n_B} = 2^{-n_B} - 2^{-(n_A+n_B)}$ | 1 | 1 |
| 1 | 1 | $(1 - 2^{-n_A}) \times (1 - 2^{-n_B}) = 1 - \left[2^{-n_A} + 2^{-n_B} - 2^{-(n_A+n_B)}\right]$ | 1 | 1/2 |

but it must be calculated numerically for more complicated distributions of skill. Table S8 provides a complete accounting of the four skill configurations for this general case, their likelihoods, and the probability of red winning conditioned on a particular configuration.

Following the calculation for the base case, the total probability that $A$ (wearing red) wins the $AB$ bout is given by

$$\Pr(\text{red wins}) = \frac{1}{2}\left(1 - 2^{-n_A} + 2^{-n_B}\right) \quad . \tag{1}$$

Direction and magnitude of the bias are determined by the relative decay rates of the two non-trivial terms (which we call "A" and "B", to reflect the sub-trees they come from). These terms represent exactly the probability that a contestant with lower skill $\pi = \epsilon$ emerges as the winner of the corresponding sub-tree.

When $n_A = n_B$, the A and B terms in Eq. (1) cancel, leading to the symmetric outcome of $\Pr(\text{red wins}) = \frac{1}{2}$. This result recovers and generalizes the result of the base case. Now suppose that we place a set of byes in the $B$ sub-tree, so that $n_B = \frac{1}{2}n_A$ (e.g., a tournament with six contestants and two byes; Section S2). The decay rate of the B term is reduced by a factor of 2, and thus for any choice of $n_A > 1$ the probability of a win by red is strictly greater than $\frac{1}{2}$. Similarly, if we place the byes in the $A$ sub-tree instead, so that $n_A = \frac{1}{2}n_B$, then the situation is reversed, and the probability of a win by red is strictly less than $\frac{1}{2}$.

##### S5.2.2 Small tournaments

We now extend the analysis to derive exact upper bounds on the bias in small tournaments. The calculated values are upper bounds because the toy model under study maximizes the variance in the skill level of contestants in the outermost round (Section S5.2.1).

From Section S2 recall that, for a tournament with $n \geq 2$ contestants, the tree includes $n_{\text{rounds}} = \lceil \log_2 n \rceil$ rounds, with $n_{\text{byes}} = 2^{n_{\text{rounds}}} - n$ byes in the outermost round, where $2^{n_{\text{rounds}}}$ is the number of contestants required to fill the round. The number of bouts across all rounds is $n - 1$. For this analysis we focus on trees with $n_{\text{rounds}} = 3$ rounds, so that $2^{n_{\text{rounds}}} = 8$ contestants are required to fill the outermost round, and we vary $n$ exhaustively. Any byes are stacked in the lower sub-tree. As above, the winners of the top and bottom sub-trees feeding into a bout wear, respectively, red and blue in the bout.

Consistent with the results of the general case (Section S5.2.1), we find bias towards wins by red in incomplete tournaments. Were we to reverse either (i) the placement of byes (i.e., byes are stacked in the upper sub-tree), or (ii) the color assignment procedure (i.e., the winner of the top sub-tree wears blue), then the direction of the bias would also reverse.

Generally, across trees of a given depth the bias is largest when the number of byes is greatest (i.e., for fewer contestants), and it tends to decrease as the number of byes decreases towards zero. Furthermore, the bias for odd numbers of byes is larger than the bias at adjacent even numbers of byes. This oscillatory behavior in the fraction of wins by red can be explained as follows. If the tree includes an even number of byes, then any sub-tree within it includes an even number of contestants. This increases the proportion of complete sub-trees within the tree, relative to trees with adjacent odd numbers of byes. The fraction of wins by red for all bouts within a complete sub-tree is always $\frac{1}{2}$ (Section S5.2.1), which thus lowers the overall fraction of wins by red compared to an incomplete sub-tree.

**$n = 5$ contestants** A tournament with five contestants includes four bouts, three byes (Figure S2a). The resulting tree can be decomposed into three components: (i) a base case of $n_A = n_B = 1$ in the lower sub-tree, the winner of which we label $Y$, (ii) a general case of $n_A = 2$ and $n_B = 1$ in the upper sub-tree, the winner of which we label $X$, and (iii) a bout that links the outcomes of these two cases together, the winner of which we label $Z$.

The expected fraction of wins by red in this tournament is the average probability across the four bouts $W, X, Y, Z$. The probability that red wins in bouts $W, X, Y$ can be calculated from Eq. (1), yielding $\frac{1}{2}$, $\frac{5}{8}$, and $\frac{1}{2}$, respectively. The probability that red wins at $Z$ must be calculated directly, and this calculation's components are given in Table S9. The result is that red wins with probability $\frac{9}{16}$. Hence, the overall fraction of wins by red is

$$f_{\text{red}} = \frac{1}{4}\left(\frac{1}{2} + \frac{5}{8} + \frac{1}{2} + \frac{9}{16}\right) = \frac{35}{64} \approx 0.547 \ .$$

**$n = 6$ contestants** A tournament with six contestants includes five bouts, two byes (Figure S2b). The resulting tree is a single instance of the general case with $n_A = 4$ and $n_B = 2$. Hence, the probability that red wins in bouts $V, W, X, Y, Z$ can be calculated from Eq. (1). All of these bouts except $Z$ are symmetric cases where $n_A = n_B$, thus the probability that red wins is $\frac{1}{2}$ in each case. The probability that red wins at $Z$ is an asymmetric case, and Eq. (1) yields $\frac{19}{32}$. Hence, the overall fraction of wins by red is

$$f_{\text{red}} = \frac{1}{5}\left(\frac{1}{2} + \frac{1}{2} + \frac{1}{2} + \frac{1}{2} + \frac{19}{32}\right) = \frac{83}{160} \approx 0.519 \ .$$

**$n = 7$ contestants** A tournament with seven contestants includes six bouts, one bye (Figure S2c). The resulting tree can be decomposed into three components: (i) a general case of $n_A = 2$ and $n_B = 1$ in the lower sub-tree, (ii) a general case of $n_A = n_B = 2$ in the upper sub-tree, and (iii) a bout that links the outcomes of these two cases together.

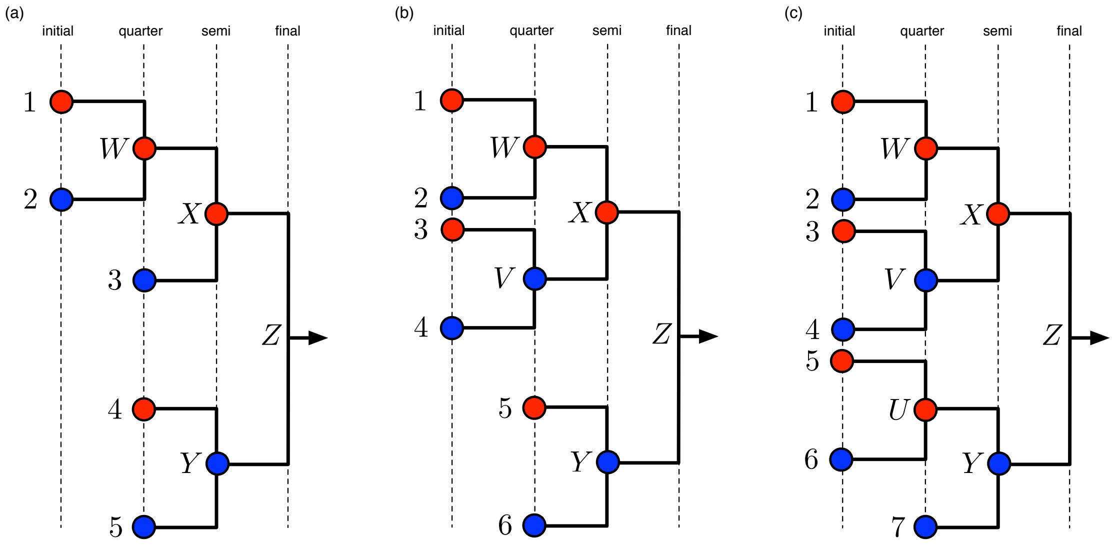

Figure S2: Tournaments including (a) $n = 5$, (b) $n = 6$, and (c) $n = 7$ contestants, with byes stacked in the lower sub-tree. The winners of the top and bottom sub-trees feeding into a bout wear, respectively, red and blue in the bout.

Table S9: Elements of calculating the outcome of bout $Z$ in the $n = 5$ tournament in Figure S2a. The probability that red (contestant at $X$) wins this bout is $\frac{36}{64} = \frac{9}{16} \approx 0.563$.

| $\pi_X$ | $\pi_Y$ | $\Pr(\pi_X, \pi_Y)$ | $\pi_{\text{winner}}$ | Pr(red wins) |
|---|---|---|---|---|
| $\epsilon$ | $\epsilon$ | $1/8 \times 1/4 = 2/64$ | $\epsilon$ | 1/2 |
| $\epsilon$ | 1 | $1/8 \times 3/4 = 6/64$ | 1 | 0 |
| 1 | $\epsilon$ | $7/8 \times 1/4 = 14/64$ | 1 | 1 |
| 1 | 1 | $7/8 \times 3/4 = 42/64$ | 1 | 1/2 |

Table S10: Elements of calculating the outcome of bout $Z$ in the $n = 7$ tournament in Figure S2c. The probability that red (contestant at $X$) wins this bout is $\frac{68}{128} = \frac{17}{32} \approx 0.531$.

| $\pi_X$ | $\pi_Y$ | $\Pr(\pi_X, \pi_Y)$ | $\pi_{\text{winner}}$ | Pr(red wins) |
|---|---|---|---|---|
| $\epsilon$ | $\epsilon$ | $1/16 \times 1/8 = 1/128$ | $\epsilon$ | 1/2 |
| $\epsilon$ | 1 | $1/16 \times 7/8 = 7/128$ | 1 | 0 |
| 1 | $\epsilon$ | $15/16 \times 1/8 = 15/128$ | 1 | 1 |
| 1 | 1 | $15/16 \times 7/8 = 105/128$ | 1 | 1/2 |

The probability that red wins in bouts $U, V, W, X, Y$ can be calculated from Eq. (1), yielding $\frac{1}{2}$, $\frac{1}{2}$, $\frac{1}{2}$, $\frac{1}{2}$, and $\frac{5}{8}$, respectively. The probability that red wins at $Z$ must be calculated directly, and this calculation's components are given in Table S10. The result is that red wins with probability $\frac{17}{32}$. Hence, the overall fraction of wins by red is

$$f_{\text{red}} = \frac{1}{6}\left(\frac{1}{2}+\frac{1}{2}+\frac{1}{2}+\frac{1}{2}+\frac{5}{8}+\frac{17}{32}\right) = \frac{101}{192} \approx 0.526 \ .$$

$n = 8$ **contestants** A tournament with eight contestants includes seven bouts, no byes, and the entire tree is a general case with $n_A = n_B = 4$. Eq. (1) yields a probability that red wins equal to $\frac{1}{2}$ in each bout in the tree. Hence, the overall fraction of wins by red is

$$f_{\text{red}} = \frac{1}{7}\left(\frac{1}{2}+\frac{1}{2}+\frac{1}{2}+\frac{1}{2}+\frac{1}{2}+\frac{1}{2}+\frac{1}{2}\right) = \frac{1}{2} \ .$$

#### S5.3 Numerical analysis

We implement a Monte Carlo simulation that can numerically calculate the null distribution for the fraction of wins by one color, conditioned on a choice of tournament and distribution of contestant skill. The simulation has a range of potential applications. For example, the resulting null distribution can be used to construct a hypothesis test for a particular distribution of contestant skill. Alternatively, it can be used to quantify the impact of different seeding procedures (e.g., seeding by skill; Section S2.5), which may introduce bias towards wins by one color over the other [2]. We leave such applications for future work, focusing instead on further investigation of systematic bias towards wins by one color over the other. Another application is in Section S6.2, where we use the simulation to determine whether any such bias applies to the 2004 and 2008 datasets.

There are multiple ways to parameterize a distribution of skill that enables a systematic investigation of how the variance in contestant skill interacts with the tournament structure. Here we assign a latent skill value $\pi$ to each contestant by drawing $\pi$ from a symmetric Beta distribution $\pi \sim \text{Beta}(\beta, \beta)$ on the unit interval, independently of the color initially assigned to the contestant. This choice of skill distribution is mathematically convenient as it guarantees that $\pi \in (0, 1)$.

We assume that a contestant's skill value is fixed over all bouts in which he participates. In the limit of $\beta \to \infty$, the Beta distribution converges on a delta function at $\pi = 0.5$, meaning that all contestants have equal skill. For finite values of $\beta$, the distribution has non-zero variance but it is symmetric about $\pi = 0.5$. When $\beta = 1$, $\pi \sim \text{Uniform}(0, 1)$, and for $\beta < 1$, the distribution exhibits a symmetric "U" shape, with the modal skill values being close to 0 or 1. This provides a one-parameter model by which we can modulate the variance in contestant skill, without making the skill distribution asymmetric around a mean of $\frac{1}{2}$. Alternative specifications of $\Pr(\pi)$, such as a Gamma distribution, yield qualitatively similar results to those presented here.

As in the mathematical analysis (Section S5.2.1), in the simulations the outcome of a bout between two contestants $r$ and $b$ is determined by the Bradley–Terry rule [20], such that $\Pr(r \text{ wins over } b) = \pi_r/(\pi_r + \pi_b)$. The winner of the bout advances to the next round. As above, the winners of the top and bottom sub-trees feeding into a bout wear, respectively, red and blue in the bout. The structure of the simulated tournaments reflects the structure of tournaments in the sports analysed (Section S2), including byes but excluding both walkovers and bouts in the consolation rounds (Section S5.1). For all simulations, results are reported using at least $10^4$ repetitions.

A first set of simulations investigates the fraction of wins by red, $f_{\text{red}}$, in tournaments of equal depth, varying (i) the number of byes, or (ii) the distribution of contestant skill. In the first scenario (Figure S3, left panels), the tree is either complete (no byes) or incomplete (including byes), and the skill distribution is such that contestants have equal skill. In the second scenario (Figure S3, right panels), the tree is complete (no byes), and the skill distribution is such that contestants have either equal or unequal skill. Across the two scenarios, selection on contestant skill is precluded in the cases where contestants have equal skill. Selection does occur in the cases where contestants have unequal skill, but there is no asymmetry in the process since the tree is complete (Section S5.1). In all cases, then, the fraction of wins by red follows a Binomial distribution centered at $f_{\text{red}} = 0.5$. It follows that, in both scenarios simulated here, a standard binomial test would be appropriate to quantify the statistical significance of an empirically observed fraction of wins by one color (Section S4.2). In line with the results of the mathematical analysis (Section S5.2), the implication is that, when the color worn by a contestant is linked to his relative position in the bout, with the assignment procedure fixed throughout the tournament, systematic bias towards wins by one color over the other arises only if both the following conditions are met: contestants have unequal skill, and the tournament tree presents "patterned" incompleteness in the outermost round due to byes. There is no bias if contestants have equal skill and/or the outermost round of the tree is complete.

Both conditions apply in a second set of simulations, which investigates the fraction of wins by red, $f_{\text{red}}$, for tournaments varying in depth and in number and placement of byes. The skill distribution is fixed with $\beta = 0.1$ (corresponding to the largest level of variance in Figs. 1**c**, **d** in the main text). The results are in Figure S4. The null distribution is centered at $f_{\text{red}} = 0.5$ for trees with a complete outermost round, but it shifts away from this value if byes are present. In agreement with the mathematical analysis (Section S5.2), the direction of the bias relative to $f_{\text{red}} = 0.5$ depends only on the placement of the byes. Specifically, if the color assignment procedure is as above, the distribution is centered at $f_{\text{red}} > 0.5$ with byes stacked in the lower sub-tree (Figure S4, top panels), and at $f_{\text{red}} < 0.5$ with byes stacked in the upper sub-tree (Figure S4, bottom panels). Overall, the magnitude of the bias decreases as the depth of the tournament increases, being largest in trees with the fewest rounds. Across trees of a given depth, the pattern is consistent with the results of the mathematical analysis (Section S5.2.2) — namely, the bias is largest when the number of byes is greatest, it tends to decrease as the number of byes decreases towards zero, and

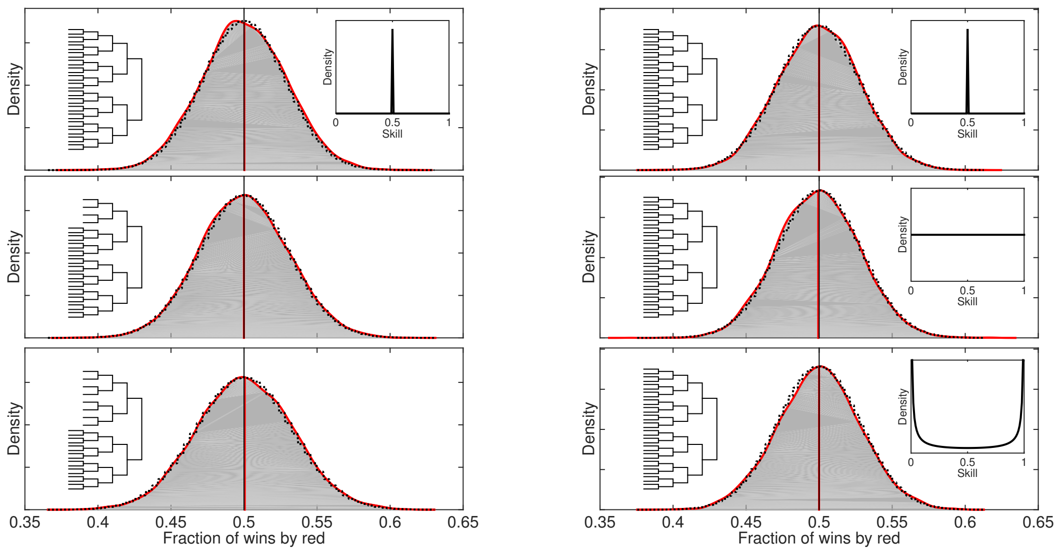

Figure S3: Simulated fraction of wins by red (solid red line) in tournaments of equal depth, varying either (i) the number of byes (left panels; trees on left), or (ii) the distribution of contestant skill (right panels; insets on right). In the first scenario the tree is either complete or incomplete (top panel vs. bottom two panels), and the skill distribution is such that contestants have equal skill. In the second scenario the tree is complete, and the skill distribution is such that contestants have either equal or unequal skill (top panel vs. bottom two panels). The dotted black line is a Binomial distribution centered at 0.5.

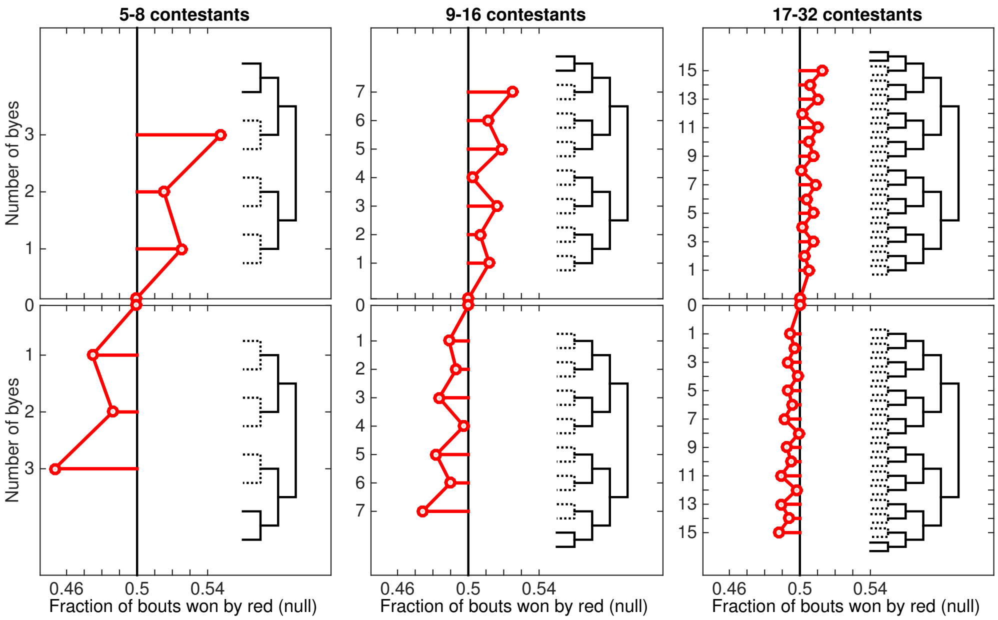

Figure S4: Simulated average fraction of wins by red for tournaments varying in depth and in number and placement of byes. Byes are stacked either in the lower sub-tree (top panels) or in the upper sub-tree (bottom panels).

it exhibits an oscillatory behavior, being larger for odd numbers of byes than for adjacent even numbers of byes. Note also that the magnitude of the bias towards wins by red with 5–8 contestants in Figure S4 matches the results of the corresponding calculations (Section S5.2.2).

#### S5.4 Summary

The tournament structure for the sports analysed here produces dependencies in the data, which may result in bias towards wins by one color over the other. This issue is not easily addressed using existing approaches in the literature and/or standard statistical procedures. To gain insight into the effect of any such bias, we presented a detailed investigation of the underlying mechanism. We build on this insight in Section S6 to investigate whether, and how, the bias affects the data for these sports at the 2004 and 2008 Olympic competitions.

We began by outlining the intuition behind the mechanism (Section S5.1). A key factor is that, in the four sports, the color worn by a contestant was linked to his relative position in the bout, with the assignment procedure fixed throughout a tournament (Section S2). In each bout, the contestant at the top of the bracket wore red in boxing and in wrestling, blue in taekwondo (Table S1).

Bias towards wins by one color over the other arises from this assignment procedure, coupled with variance in contestant skill and with incompleteness in the tournament tree. The underlying mechanism is asymmetric selection on contestant skill. Selection on contestant skill comes from the tournament itself. By definition, contestants with lower skill are more likely to be eliminated as the tournament progresses. As a result of this selective process, the distribution of contestant skill shifts progressively towards higher values.

The shift is symmetric in complete tournaments, and asymmetric in incomplete ones. In particular, "patterned" incompleteness in the tournament tree due to byes leads to weaker selection in one of the two sub-trees — specifically, in the sub-tree with more byes (and thus fewer contestants). Throughout the tournament, then, if the two sub-trees feeding into a bout include different numbers of contestants, the expected skill level of the contestant in one bout position (top vs. bottom of the bracket) will exceed his opponent's, making the contestant in that position more likely to win. If the color assignment procedure is as above, there will be more wins by contestants wearing the color associated with that position, and thus a bias towards wins by one color over the other.

Byes are a source of "patterned" incompleteness because they are always placed in the outermost round of the tree, stacked either in the upper vs. lower sub-tree (Section S2.1). For example, consider a case where, in each bout, the contestants at the top and bottom of the bracket wear red and blue, respectively, and byes are stacked in the lower sub-tree. In this case, the selection on contestant skill will be weaker in the lower sub-tree, which includes more byes (and thus fewer contestants). Throughout the tournament, if the two sub-trees feeding into a bout include different numbers of contestants, the expected skill level of the contestant at the top of the bracket will exceed his opponent's, making the contestant in that bout position more likely to win. Overall, there will tend to be more wins by contestants wearing red, and thus a bias towards wins by one color over the other.

We term this bias "systematic" to distinguish it from bias that may arise from walkovers, which we term "idiosyncratic". Like byes, walkovers are a source of incompleteness, but in this case there is no pattern in the placement of the missing bouts with respect to each other and, ultimately, with respect to color (Section S2). Thus, whether walkovers do result in asymmetric selection on contestant skill, and whether this, in turn, results in bias towards wins by one color over the other, depends on the specific configuration. It follows that direction and magnitude of the bias can be predicted based on a few key factors in the case of byes, but they are not easily predicted in the case of walkovers. It also follows that, if a tree presents both systematic and idiosyncractic biases, these may interact synergistically or antagonistically — their combined effect again depends on the specific configuration, such that direction and magnitude of the overall bias are also not easily predicted.

For simplicity, then, we focused on the effect of byes, to the exclusion of both walkovers and bouts in the consolation rounds. We explored the mechanism through mathematical analysis (Section S5.2), based on a toy model of a single-elimination tournament. The analysis was in two parts. In the first part, we demonstrated the existence of the mechanism in a general setting (Section S5.2.1). In the base case of a single bout, the tournament effectively enhances the probability that the winner of the bout is the contestant with higher skill. The tournament selects as the winner the contestant with higher skill, and the outcome of the bout is thus symmetric with respect to color. In the general case including multiple bouts, if the two sub-trees feeding into a bout include different numbers of contestants, then the expected skill level of the contestant from the larger sub-tree exceeds his opponent's, making the contestant from that sub-tree more likely to win. This is because each sub-tree exerts a selective effect on skill, and the effect grows in strength with the number of contestants in the sub-tree. If the color assignment procedure is as above, direction and magnitude of the bias both depend on the relative numbers of contestants in the two sub-trees feeding into the bout.

In the second part of the mathematical analysis, we derived upper bounds on systematic bias in small tournaments (Section S5.2.2). We showed that, across trees of a given depth (i.e., for a given number of rounds), the bias is largest when the number of byes is greatest (i.e., for fewer contestants), and it tends to decrease as the number of byes decreases towards zero. Furthermore, the bias exhibits an oscillatory behavior, being larger at odd numbers of byes than at adjacent even numbers of byes.

We then extended the investigation to more general versions of the model through numerical analysis (Section S5.3). The results of this analysis confirmed those of the mathematical analysis (Section S5.2). A first set of simulations showed that, when the color worn by a contestant is linked to his relative position in the bout, with the assignment procedure fixed throughout the tournament, systematic bias towards wins by one color over the other arises if both the following conditions are met: contestants have unequal skill, and the tournament tree presents "patterned" incompleteness in the outermost round due to byes. If contestants have equal skill and/or the outermost round is complete, the null distribution of the fraction of wins by one color follows a Binomial distribution centered at 0.5.

A second set of simulations showed that, when both conditions are met, the null distribution shifts towards wins by one color or the other. For a given color assignment procedure, the direction of the bias depends only on the placement of byes in the outermost round. Specifically, the bias is towards the color associated with the top bout position if byes are stacked in the lower sub-tree, and towards the color associated with the bottom bout position if byes are stacked in the upper sub-tree. The magnitude of the bias decreases as the depth of the tournament increases, being largest in trees with the fewest rounds. Across trees of a given depth, the behavior is as predicted by the mathematical analysis (Section S5.2.2).

### S6 Triangulation

We present a final set of analyses to evaluate Hill & Barton's [1] claim of an effect of red on the outcomes of Olympic combat sports. Using data relating to the two Olympic competitions for the four sports analysed here (Section S3), we triangulate insights we obtained from description of the tournament structure and related aspects for the sports (Section S2), from the replication analysis (Section S4), and from investigation of the mechanism leading to bias towards wins by one color over the other (Section S5).

We begin by outlining qualitative considerations about each case (Section S6.1). Next, we use the Monte Carlo simulation described in Section S5.3 to determine whether the data present bias towards wins by one color over the other (Section S6.2). Finally, we use the analytical approach described in Section S4.2 to test the prediction of a majority of wins by red in sub-sets of the data, so as to minimize the effect of the bias (Section S6.3).

#### S6.1 Qualitative considerations

As discussed in Section S5, in a single-elimination tournament bias towards wins by one color over the other may arise from missing bouts due to byes and/or to walkovers. We present separately considerations that apply to the main tournament trees (Section S6.1.1) and to the consolation rounds (Section S6.1.2).

##### S6.1.1 Main tournament trees

We discussed in Section S5.1 that byes are a source of "patterned" incompleteness in a tournament tree: they are always placed in the outermost round, stacked either in the upper sub-tree or in the lower one (Section S2). Coupled with variance in contestant skill, they result in systematic bias towards wins by one color over the other if the color worn by a contestant is linked to his relative position in the bout, with the assignment procedure fixed throughout the tournament. The bias is systematic because its direction and magnitude can be predicted based on a few key factors — namely, the distribution of contestant skill, the color assignment procedure, the placement of the byes, and their number relative to the number of contestants entering the tournament (Section S5.1).

Relevant features of the tournament structure and related aspects for each sport, by year, are summarized in Table S1, including the average value of the completeness fraction linked to byes, $\rho$, across weight classes. Recall from Section S2.1 that $\rho = 1.00$ for a tournament with no byes, and $\rho < 1.00$ otherwise. Thus, the average value of $\rho$ across weight classes provides a simple measure of how systematically (in)complete the outermost round of the tournament tended to be for a sport in a given year.

The color assignment procedure is as above in all four of the sports, with the contestant at the top/bottom of the bracket wearing red/blue in boxing and in wrestling, and blue/red in taekwondo (Table S1). We can assume variance in skill to apply in each case: although the true variance is not known (Section S4.1.2), it seems unlikely that all contestants entering an

Olympic tournament would have equal skill. Crucially, however, "patterned" incompleteness in the main tournament tree applies in boxing and wrestling only, as reflected in the average values of $\rho$ across weight classes (Table S1). Recall from Section S2.3 that in taekwondo the competition for each weight class was arranged as a single-elimination tournament with $n = 16$ contestants. Thus, in each weight class the outermost round was a complete round of 16 (eighth-finals), and there were no byes.

Consequently, for the main tournament trees we expect systematic bias towards wins by one color over the other only in boxing and in wrestling. We can also formulate specific predictions about the direction of the bias. In Section S5 we showed that, for a given color assignment procedure, the direction depends only on the placement of the byes in the outermost round. Specifically, if byes are stacked in the lower/upper sub-tree, then the bias is towards the color associated with the top/bottom bout position (i.e., red/blue in boxing and in wrestling; Table S1). Byes were stacked in the lower sub-tree in 2004 wrestling, and in the upper sub-tree in 2008 wrestling and in boxing (Table S1). Therefore, in the main tournament trees we expect systematic bias towards wins by red in 2004 wrestling, and towards wins by blue in 2008 wrestling and in boxing.

Generally, for a given distribution of contestant skill we expect the magnitude of the bias to reflect the depth of the tournament (i.e., the number of rounds in the tree), coupled with the relative numbers of byes vs. contestants in the outermost round (Section S5.4). In the absence of information about the true variance in skill, we cannot formulate specific predictions about the absolute magnitude of the bias in each case. Still, assuming comparable distributions of skill for the two sports and across the two years, we can generate broad qualitative expectations in relative terms. The tournament trees included three rounds in 2004 wrestling (i.e., an outermost round of 8), five in 2008 wrestling (i.e., an outermost round of 32), and four or five in boxing (i.e., an outermost round of 16 or 32, respectively; Table S1). Therefore, on account of the relative depth of the tournaments, we expect the bias to be larger in 2004 wrestling than in 2008 wrestling and in boxing. Recall from Section S2.2.1 that the boxing tournaments with an outermost round of 16 always included $n = 16$ contestants, thus they were always complete. In the other boxing tournaments $n$ ranged from 27 to 29 contestants in an outermost round of 32 (Section S2.2.1). For comparison, in the 2008 wrestling tournaments $n$ ranged from 19 to 21 contestants in an outermost round of 32 (Section S2.4.1). Therefore, on account of the relative number of byes vs. contestants, as reflected in the average values of $\rho$ across weight classes (Table S1), we expect the bias to be larger in 2008 wrestling than in boxing.

We can now turn to the effect of walkovers in the main tournament trees. As discussed in Sections S5.1 and S5.4, if the color assignment procedure is as above, and contestants vary in skill, walkovers may also result in bias towards wins by one color over the other — whether they do depends on the specific configuration. The reason is that walkovers can occur anywhere on the tree. Therefore, there is no pattern in the placement of the missing bouts with respect to each other and, ultimately, with respect to color. In this case the bias is idiosyncratic, in that its direction and magnitude are not easily predicted based on relevant features of the tournament structure and related aspects. Furthermore, if a tree includes both byes and walkovers, their combined effect also depends on the specific configuration: the systematic bias arising from byes may interact synergistically or antagonistically with any idiosyncratic bias arising from walkovers. Consequently, direction and magnitude of the overall bias are also not easily predicted.

It follows that we can generate only broad qualitative expectations about walkovers, relating specifically to the likely effect on the overall bias. Neither taekwondo nor 2004 wrestling featured walkovers in the main tournament trees. In boxing and in 2008 wrestling the number of walkovers in the main tournament trees was small in relation to the size of the tournaments. Specifically, there were five walkovers out of 272 bouts in 2004 boxing, two out of 272 bouts in 2008 boxing, and one out of 132 bouts in 2008 free-style wrestling. Therefore, we expect no idiosyncratic bias in taekwondo and in 2004 wrestling, and any that may arise in boxing and in 2008 wrestling is likely to be small. Depending on its direction, it may modulate the magnitude of the systematic bias due to byes, by dampening or enhancing it. However, it is unlikely to do so to the extent that the direction of the overall bias would actually reverse, relative to the systematic bias.

In sum, for the main tournament trees we expect no bias in taekwondo, and an overall bias towards wins by red in 2004 wrestling and towards wins by blue in 2008 wrestling and in boxing. Furthermore, assuming the effect of walkovers to be small and comparable across cases, we expect the overall bias to be larger in 2004 wrestling than in 2008 wrestling, and larger in 2008 wrestling than in boxing.

###### S6.1.2 Consolation rounds

Across the four sports, consolation rounds are found in taekwondo (i.e., the repechage contests; Section S2.3.3) and in wrestling (i.e., the 3–4 and 5–6 final rounds in 2004, and the repechage contests in 2008; Section S2.4.3). It is useful to think of each of these rounds as a separate “ancillary” tournament, featuring the same color assignment procedure as the corresponding main tournament. In each ancillary tournament, missing bouts due to byes and/or walkovers may induce asymmetric selection on contestant skill, which in turn may induce bias towards wins by one color over the other, with the overall effect depending on the specific configuration. Therefore, the consolation rounds are also a potential source of bias in the data as a whole.

Any bias that may arise in the ancillary tournaments is idiosyncratic, on account of how these tournaments were populated. In particular, the distribution of skill cannot be assumed to be randomized in the way it was in the in the outermost round of the main tournaments. Recall from Section S2.5 that in the main tournaments both the position of contestants and the placement of byes were independent of contestant skill: initial seeding was based on a random draw, and byes were stacked in either the upper sub-tree or the lower one. By contrast, in the ancillary tournaments the position of contestants was linked to performance in the main tournaments, and byes were placed to reflect progression in those. For example, in all cases the losers of the semi-final rounds in the main tournaments competed for bronze in the consolation rounds. Specifically, they competed in the 3–4 final round in 2004 wrestling, and they received one bye to the bronze round(s) in taekwondo and the equivalent of two byes to the bronze rounds in 2008 wrestling. Assuming that performance in the main tournaments was linked to contestant skill, the position of contestants and the placement of byes in the ancillary tournaments was thus not independent of skill. In particular, contestants entering the ancillary tournaments may have been subjected to different levels of selection in the main tournaments, potentially leading to bias towards wins by one color over the other (Section S5).

Whether any such bias applies in each case depends on relevant features of both main and ancillary tournaments, as we discuss below with reference to the bronze round(s). Furthermore, the bias may interact synergistically or antagonistically with any bias arising from walkovers in the ancillary tournaments. In 2008 wrestling the were no walkovers in the consolation rounds. In taekwondo and in 2004 wrestling the number of walkovers in the consolation rounds was comparable to the number of walkovers in the main tournament trees (Section S6.1.1), against a considerably smaller number of bouts. Therefore, the number of walkovers was large in relation to the size of the tournaments. Specifically, there were five walkovers out of 20 bouts in 2004 taekwondo, one out of 16 bouts in 2008 taekwondo, three out of 13 bouts in 2004 Greco-Roman wrestling, and three out of 14 bouts in 2004 free-style wrestling.

To illustrate the effect of all these factors, combined, we focus on the outcome of the bronze round(s) in each case. In taekwondo the ancillary tournaments were symmetric with respect to color (Section S2.3.3). All walkovers occurred "earlier" in the repechage contests than the bronze rounds, thus they potentially had an effect on these rounds. The main tournaments featured neither walkovers (Section S6.1.1) nor byes (Section S2.5). Therefore, any two contestants in "equivalent" positions in the repechage contest(s) for each weight class had likely experienced comparable levels of selection in the main tournament tree. Consequently, we expect no bias towards wins by one color in the outcome of the bronze round(s), except any arising from the specific configuration of walkovers in the repechage contest(s) (Section S5).

In 2004 wrestling the ancillary tournaments were also symmetric with respect to color (Section S2.4.3). All walkovers occurred in the 5–6 final rounds, thus they had no effect on the 3–4 final rounds. The main tournaments featured no walkovers (Section S6.1.1), and byes were stacked in the lower sub-tree (Section S2.5). Therefore, the two contestants competing for bronze in the 3–4 final round for each weight class had likely experienced different levels of selection in the main tournament tree. Specifically, we expect stronger selection on the contestant entering from the upper sub-tree, leading to bias towards wins by red in the outcome of the bronze round (Section S5).

Finally, in 2008 wrestling the ancillary tournaments were asymmetric with respect to color (Section S2.4.3), and there were no walkovers in the consolation rounds. With one exception (namely, one walkover in the eighth-final round for the 96kg weight class in free-style wrestling), the main tournaments featured no walkovers (Section S6.1.1), and byes were stacked in the upper sub-tree (Section S2.5). Therefore, any two contestants in "equivalent" positions in the repechage contests for each weight class had likely experienced different levels of selection in the main tournament tree. Specifically, we expect weaker selection on the contestant byed to the bronze round of the contest corresponding to the upper half of the main tournament tree, compared to his counterpart in the other contest. Consequently, we expect stronger bias towards wins by blue in the outcome of the bronze round of the contest corresponding to the lower half of the tree. The effect across the two contests is contingent on the position of additional byes included in the consolation rounds on an *ad-hoc* basis (Section S2.4.3).

In sum, for the ancillary tournaments we cannot formulate even broad expectations about whether overall bias towards wins by one color over the other obtains in each case.

#### S6.2 Numerical analysis

In Section S6.1 we outlined potential sources of bias in the data towards wins by one color over the other. We now turn to the Monte Carlo simulation described in Section S5.3 to determine their overall effect. Specifically, we numerically calculate the distribution of the fraction of wins by red, $f_{\text{red}}$, under the null hypothesis (no effect of red) for (i) the observed tournaments, and (ii) equivalent tournaments with no missing bouts. In practice, we implement two sets of simulations for each weight class, varying the distribution of contestant skill within each set.

The first set of simulations is parametrized by the tournament for the weight class, exactly as observed. In particular, the simulated tournaments include any consolation rounds and any walkovers that occurred in the observed tournaments; recall that both were excluded from the simulations implemented in Section S5.3. In each simulated tournament, then, any consolation rounds follow the rules that applied in the observed tournament. Furthermore, if the observed tournament included a win by walkover, the relevant contestant automatically advances to the next round in the simulated tournament.

The second set of simulations is parametrized by an equivalent tournament with no missing bouts. To this end, we first "complete" the tournament tree by removing any missing bouts due to byes and/or to walkovers. This step necessarily adds bouts to the simulated tournament compared to the observed tournament, which increases the sample size and thus decreases the variance of the null distribution of $f_{\text{red}}$. To correct this difference, we then prune a corresponding number of bouts from locations of symmetry in the tree (e.g., the final round in any sport, the 3–4 or 5–6 final rounds in 2004 wrestling, or an entire repechage contest in taekwondo and in 2008 wrestling; Section S2).

We showed in Section S5.3 that if contestants have equal skill and/or the tournament tree is complete, then the null distribution of $f_{\text{red}}$ follows a Binomial distribution centered at 0.5. Having removed any missing bouts from the simulated tournaments, in the second set of simulations we expect the distribution of $f_{\text{red}}$ to remain centered at 0.5 for any distribution of contestant skill. Therefore, deviations from this pattern in the first set of simulations can be taken as evidence of bias towards wins by one color over the other, induced by the missing bouts in the observed tournaments.

For all simulations, results are reported using $10^5$ repetitions, combined over all weight classes, for the four sports individually and aggregated over boxing and taekwondo, by year (Figures S5 and S6). The results aggregated over all sports by year are in Figs. 1**c, d** in the main text. Each panel shows the null distributions of $f_{\mathrm{red}}$ for the two sets of simulations as a function of variance in contestant skill, together with the observed value of $f_{\mathrm{red}}$.

In each sport individually by year (Figures S5 and S6), as expected there is no bias with no variance in skill, and as the variance in skill increases, so does the magnitude of the bias. The increase is more substantial in wrestling than in boxing and in taekwondo, and it is larger in 2004 wrestling than in 2008 wrestling. Conversely, in boxing and in taekwondo the increase is comparable across years, with minimal bias even at the highest levels of variance in skill. The direction of the bias is towards wins by blue in boxing and in 2008 wrestling, and towards wins by red in taekwondo and in 2004 wrestling.

This pattern is fully in line with the qualitative considerations outlined in Section S6.1. In boxing there were no consolation rounds, thus the overall bias results from missing bouts due to byes and/or to walkovers in the main tournament trees. The magnitude of the bias reflects the small number of missing bouts relative to the size of the tournaments (Section S6.1.1).

In taekwondo there were no missing bouts in the main tournament trees (Section S6.1.1). Given that the ancillary tournaments were symmetric with respect to color, the overall bias can be attributed to walkovers in the repechage contest(s) (Section S6.1.2). The magnitude of the bias likely reflects the complete symmetry of both main and ancillary tournaments, balanced against the relatively large number of walkovers (Section S6.1).

As for wrestling, the shift in direction of the overall bias reflects the substantial changes in tournament structure that were implemented between the two competitions. Crucially, the changes included a shift in the placement of byes from the lower sub-tree in 2004 to the upper one in 2008 (Section S2.4). The color assignment procedure did not change, however: the contestant in the top/bottom bout position wore red/blue in both years (Table S1). Recall from Section S5 that, for a given color assignment procedure, the direction of the bias depends only on the placement of the byes in the outermost round. As expected, then, the bias is towards the color associated with the top bout position in 2004, and towards the color associated with the bottom bout position in 2008 (Section S6.1.1). The magnitude of the bias reflects the considerable asymmetry of both main and ancillary tournaments, arising from the relatively large number of missing bouts due to byes and to walkovers (Section S6.1).

Across the four sports by year (Figs. 1**c, d** in the main text), again there is no bias with no variance in skill, and as the variance in skill increases, so does the magnitude of the bias. The increase is comparable across years — even though, as noted above, it is larger in 2004 wrestling than in 2008 wrestling, and comparable across years in boxing

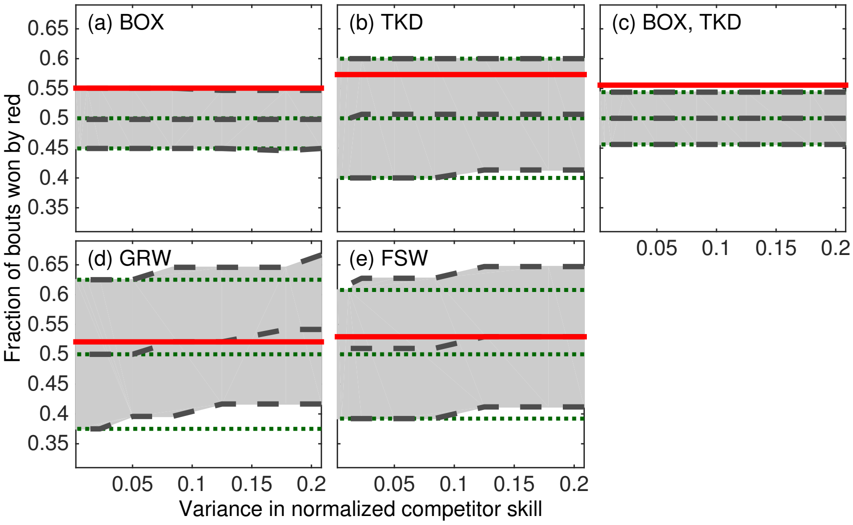

Figure S5: Quantiles (from top to bottom, 95%, 50%, and 5%) for the distribution of the fraction of wins by red, $f_{\mathrm{red}}$, under the null hypothesis on (i) the observed tournaments at the 2004 Athens Olympics (dashed grey lines, grey fill), and (ii) equivalent tournaments with no missing bouts (dotted green lines), for (a) boxing (BOX), (b) taekwondo (TKD), (c) aggregated over boxing and taekwondo (BOX, TKD), (d) Greco-Roman wrestling (GRW), and (e) free-style wrestling (FSW). The solid red line shows the observed value of $f_{\mathrm{red}}$.

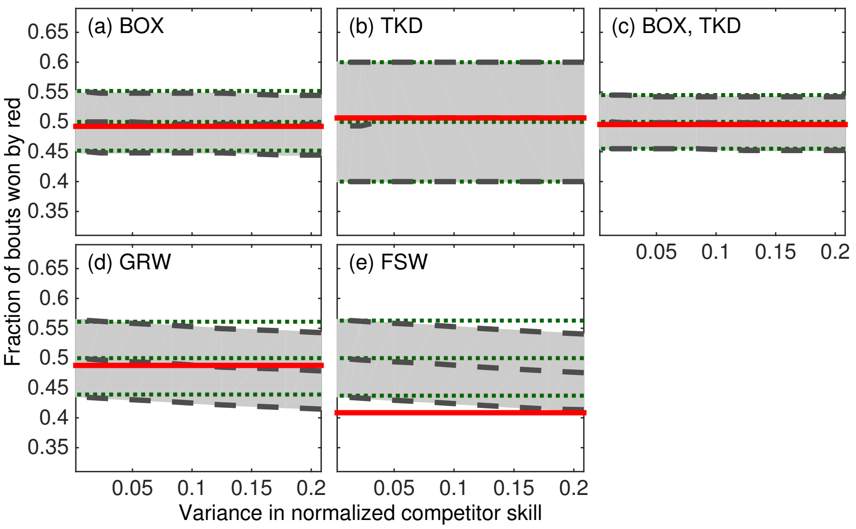

Figure S6: Quantiles (from top to bottom, 95%, 50%, and 5%) for the distribution of the fraction of wins by red, $f_{\text{red}}$, under the null hypothesis on (i) the observed tournaments at the 2008 Beijing Olympics (dashed grey lines, grey fill), and (ii) equivalent tournaments with no missing bouts (dotted green lines), for (a) boxing (BOX), (b) taekwondo (TKD), (c) aggregated over boxing and taekwondo (BOX, TKD), (d) Greco-Roman wrestling (GRW), and (e) free-style wrestling (FSW). The solid red line shows the observed value of $f_{\text{red}}$.

and in taekwondo (Figures S5 and S6). This pattern reflects variation in the number of bouts available for analysis in 2004 compared to 2008. In particular, the 2008 dataset more than trebles the number of bouts in both Greco-Roman and free-style wrestling relative to the 2004 dataset (Section S2.4.4). The number of bouts is comparable across years in the other sports (Section S3); therefore, wrestling accounts for a higher proportion of bouts in the 2008 dataset than in the 2004 dataset. This difference in proportions between years effectively "offsets" the larger bias in 2004 wrestling compared to 2008 wrestling.

In both years the magnitude of the bias is minimal even at the highest levels of variance in skill (Figs. 1**c, d** in the main text), being comparable to the overall bias in boxing and in taekwondo (Figures S5 and S6). The direction of the bias is towards wins by red in 2004 and towards wins by blue in 2008, consistent with the shift in direction of the overall bias between the two competitions in wrestling (Figures S5 and S6). Exclusion of the corresponding simulation runs confirms that wrestling is the primary source of bias in the results aggregated over all sports (Figures S5 and S6).

These findings indicate that, for data relating to wrestling, a correctly parameterized test of Hill & Barton's [1] hypothesis cannot be constructed without knowledge of the variance in contestant skill. Absent this information, a conservative approach is to exclude the corresponding data from further analysis.

#### S6.3 Data analysis

In Section S6.2 we showed that data relating to the two Olympic competitions for the four sports analysed here present bias towards wins by one color over the other. In this section we conduct a final analysis of Hill & Barton's [1] claim of an effect of red on the outcomes of Olympic combat sports. We restrict the analysis to data for boxing and for taekwondo, as the bias is most prominent in wrestling (Section S6.2). Furthermore, we focus on the fraction of bouts won by red, to eschew problematic assumptions about the distribution of rounds, weight classes, and points scored (Sections S4.1.1 and S4.1.2). We apply the analytical approach developed in Section S4.2, which specifies the tests in line with Hill & Barton's [1] hypothesis, in addition to providing adjustment for multiple hypothesis testing. For ease of comparison with the analysis underpinning Hill & Barton's [1] main claim (Section S4.1.1), we analyse the data for each sport individually by year and aggregated in all possible sport/year combinations.

Specifically, we use a series of one-sided binomial tests ($H_0 : f_{\text{red}} \leq 0.5; H_{\text{A}} : f_{\text{red}} > 0.5$; $\alpha = 0.05$) to test for an effect in the fraction of bouts won by red, separately for boxing and for taekwondo and aggregated over the two sports, both by year and aggregated over the two years. This gives a total of nine tests, summarized in Table S11. In addition to the raw $p$-values, the table includes the $100(1 - 2\alpha)\% = 90\%$ confidence intervals for $f_{\text{red}}$, calculated using Wilson's [9] method [10], and the $p$-values adjusted for multiple hypothesis testing using four methods (Bonferroni [11, 12], Holm [13], Benjamini & Hochberg [14], and Benjamini & Yekutieli [15]).

Focusing on the raw $p$-values, only one result is significant at the $\alpha = 0.05$ level, relating to the data aggregated over the two sports in 2004. The other eight tests provide no evidence to support the prediction of a majority of red wins, not even those relating to either sport individually in 2004 (Table S11).

In theory, it is possible for the "true" effect of red to be masked by bias towards wins by blue, as expected in the boxing data in both years and in the data aggregated over the two sports in 2008 (recall that the magnitude of the bias is minimal even at the highest levels of variance in contestant skill; Section S6.2). Yet the boxing data return qualitatively different patterns for the two years, with $f_{\text{red}} > 0.5$ in 2004 and $f_{\text{red}} < 0.5$ in 2008 (Figures S5 and S6). This difference is unlikely to arise from substantially stronger bias towards wins by blue in the 2008 competition: the size of the tournaments is comparable between years, as is the number of missing bouts due to byes and to walkovers (Section S6.1.1). It follows that the small bias towards wins by red observed in the 2004 boxing data is likely due to chance. This interpretation is corroborated by the pattern in the boxing data aggregated over the two years, and by inspection of the relevant confidence intervals (Table S11).

Returning to the full set of results, then, rejection of the null hypothesis for the 2004 data aggregated over the two sports may itself be an artifact, driven by the boxing data. Consistently, adjustment for multiple hypothesis testing indicates that this rejection is likely spurious (Table S11). Specifically, controlling the family-wise error rate at level $\alpha = 0.05$ — that is, evaluating the adjusted $p$-values against this threshold — produces no statistically significant results for either method (i.e., Bonferroni [11, 12], Holm [13]). The adjusted $p$-values indicate that a family-wise error rate $\alpha \geq 0.20$ would be required to reject the null hypothesis rejected with no adjustment. This value corresponds to a high probability of committing a Type I error (rejecting the null hypothesis, when the null hypothesis is in fact true — i.e., a false positive; Section S4.2).

Similarly, controlling the false discovery rate at 5%, 10%, or 15% — that is, evaluating the adjusted $p$-values against these thresholds — produces no statistically significant results for either method (i.e., Benjamini & Hochberg [14], Benjamini & Yekutieli [15]). With regards to the null hypothesis rejected with no adjustment, the adjusted $p$-values indicate that a false discovery rate of at least 20% (under Benjamini & Hochberg [14]) or 58% (under Benjamini & Yekutieli [15]) would be required for that null hypothesis to be included in the set of rejections. Both values correspond to high "tolerance" for false rejections (Section S4.2).

Evaluated as a whole, these results refute Hill & Barton's [1] claim of an effect of red on the outcomes of Olympic combat sports. There is no evidence of such an effect in data for the 2004 and 2008 Olympic competitions.

Table S11: Results of one-sided binomial tests in data for boxing (BOX), taekwondo (TKD), and aggregated over the two sports (BOX, TKD), at the 2004 Athens Olympics and the 2008 Beijing Olympics. The tests compare the number of bouts won by red, $n_{\mathrm{red}}$, to the total number of bouts, $n_{\mathrm{tot}}$, with $f_{\mathrm{red}} = n_{\mathrm{red}}/n_{\mathrm{tot}}$. Included are the raw $p$-values ($H_0 : f_{\mathrm{red}} \leq 0.5; H_{\mathrm{A}} : f_{\mathrm{red}} > 0.5$; $\alpha = 0.05$), the 90% confidence intervals for $f_{\mathrm{red}}$, and the $p$-values adjusted for multiple hypothesis testing using four methods (B: Bonferroni [11, 12]; H: Holm [13]; BH: Benjamini & Hochberg [14]; BY: Benjamini & Yekutieli [15]).

| Year | Sport(s) | $n_{\mathrm{red}}$ | $n_{\mathrm{tot}}$ | $f_{\mathrm{red}}$ | 90% CI | $p$ | | | | |
|---|---|---|---|---|---|---|---|---|---|---|
| | | | | | | Raw | B | H | BH | BY |
| 2004 | BOX | 147 | 267 | 0.551 | 0.500–0.600 | 0.056 | 0.501 | 0.446 | 0.251 | 0.709 |
| 2004 | TKD | 43 | 75 | 0.573 | 0.478–0.663 | 0.124 | 1.000 | 0.744 | 0.277 | 0.783 |
| 2004 | BOX, TKD | 190 | 342 | 0.556 | 0.511–0.599 | 0.023 | 0.204 | 0.204 | 0.204 | 0.576 |
| 2008 | BOX | 133 | 270 | 0.493 | 0.443–0.542 | 0.620 | 1.000 | 1.000 | 0.620 | 1.000 |
| 2008 | TKD | 38 | 75 | 0.507 | 0.413–0.600 | 0.500 | 1.000 | 1.000 | 0.620 | 1.000 |
| 2008 | BOX, TKD | 171 | 345 | 0.496 | 0.452–0.540 | 0.585 | 1.000 | 1.000 | 0.620 | 1.000 |
| Both | BOX | 280 | 537 | 0.521 | 0.486–0.557 | 0.171 | 1.000 | 0.856 | 0.277 | 0.783 |
| Both | TKD | 81 | 150 | 0.540 | 0.473–0.606 | 0.185 | 1.000 | 0.856 | 0.277 | 0.783 |
| Both | BOX, TKD | 361 | 687 | 0.525 | 0.494–0.557 | 0.097 | 0.875 | 0.681 | 0.277 | 0.783 |

### S7 Discussion

We provided multiple lines of analysis, together with new data, to evaluate Hill & Barton's [1] hypothesis of an effect of red on the outcome of human competitive interactions. The results reported in [1] were based on analysis of data for four combat sports at the 2004 Athens Olympics.

First, we showed that the results underpinning the claim of an effect of red in these data are not robust. Consistently, we found that the pattern does not hold in equivalent data for the 2008 Beijing Olympics, even replicating the analytical approach used in [1].

Second, we uncovered several shortcomings with the research design, analysis, and interpretation underlying the results reported in [1], from issues of test mis-specification to issues with interpretation. Therefore, we devised an alternative analytical approach, and again we showed that there is no evidence of an effect of red in either dataset.

Third, we investigated mathematically and numerically the issue of dependencies in the data linked to the tournament structure for the sports analysed. We demonstrated that asymmetric selection on contestant skill in a tournament can result in bias towards wins by one color over the other.

Fourth, we investigated via simulation whether any such bias obtains in the two datasets. We found bias towards wins by red in the outcomes of the 2004 competition, and towards wins by blue in the outcomes of the 2008 competition.

Fifth, we conducted a final analysis of the data that minimized the effect of the bias, and again we showed that there is no evidence of an effect of red in either dataset.

These findings converge to indicate that the effect of red on human competition reported by Hill & Barton [1] is likely spurious. Previous claims attributed the pattern in [1] to random variation [e.g., 21]; our work shows instead that it may reflect bias towards wins by red in the outcomes of the 2004 competition, linked to the tournament structure. Therefore, it is not necessary to invoke any of the behavioral, structural, or other confounds that have been proposed over the years as alternatives to the original interpretation in [1], such as differences in visibility between colors, asymmetries in prior experience across contestants (e.g., win–lose effects and/or number of previous bouts fought), and differences in recovery time due to variation in intervals between contests [2, 18, 19]. Of course, based on our results we cannot exclude that these or other factors may operate in the sports analysed [see e.g., 22, for suggestive evidence that competition judges in taekwondo favor contestants wearing red]. Yet in the absence of evidence that such factors did indeed apply to competitions for these sports at the 2004 Athens Olympics, our explanation provides the most parsimonious and comprehensive account to date of the pattern reported in [1].

Hill & Barton [1] framed their results as evidence that the effect of red on human competition is a response shaped by sexual selection, analogous to the response observed in other animal species. A large body of work has developed since, straddling biology and the social sciences, building on Hill & Barton's [1] influential study [reviewed in 23–25]. Given the difficulties that arise in seeking to disentangle a "real" effect of red from potential confounds in observational data, researchers have increasingly turned to experiments to demonstrate the existence of such an effect. Irrespective of the approach used, work in this area routinely points to the results in [1] as the key piece of evidence that the effect of red on human competition is a response shaped by sexual selection and, by implication, as key evidence of “parallels between the human and nonhuman response to color” [25, p. 115]. Our refutation of the results in [1] calls for a critical re-evaluation of this body of work, together with a re-assessment of the theoretical premise of future studies that depend on it conceptually. In particular, caution is required when invoking an evolutionary basis for any effect of red on human competition, and on human behavior more generally, based on the results reported in [1].

To be clear, we do not dispute the notion that the human response to color has been shaped by evolutionary processes over time — this seems incontrovertible. Yet in what way it has been shaped by them, and how this is reflected in present-day human behavior, remain open questions.

### References

1. Hill, R. A. & Barton, R. A. Red enhances human performance in contests. *Nature* **435,** 293–293 (2005).
2. Dijkstra, P. D. & Preenen, P. T. No effect of blue on winning contests in judo. *Proceedings of the Royal Society of London B: Biological Sciences* **275,** 1157–1162 (2008).
3. Seife, C. *Red does not enhance human performance in the Olympics* `http://www.users.cloud9.net/~cgseife/SeifeOlympicsManuscript12February.pdf`. [Online; retrieved on 2014-03-06].
4. Simmons, J. P., Nelson, L. D. & Simonsohn, U. False-positive psychology: undisclosed flexibility in data collection and analysis allows presenting anything as significant. *Psychological Science* **22,** 1359–1366 (2011).
5. Gelman, A. & Loken, E. The statistical crisis in science. *American Scientist* **102,** 460–465 (2014).
6. Peng, R. D. Reproducible research in computational science. *Science* **334,** 1226–1227 (2011).
7. Shaffer, J. P. Multiple hypothesis testing. *Annual Review of Psychology* **46,** 561–584 (1995).
8. Gelman, A. & Stern, H. The difference between "significant" and "not significant" is not itself statistically significant. *The American Statistician* **60,** 328–331 (2006).
9. Wilson, E. B. Probable inference, the law of succession, and statistical inference. *Journal of the American Statistical Association* **22,** 209–212 (1927).
10. Agresti, A. & Coull, B. A. Approximate is better than "exact" for interval estimation of binomial proportions. *The American Statistician* **52,** 119–126 (1998).
11. Dunn, O. J. Estimation of the medians for dependent variables. *Annals of Mathematical Statistics* **30,** 192–197 (1959).
12. Dunn, O. J. Multiple comparisons among means. *Journal of the American Statistical Association* **56,** 52–64 (1961).
13. Holm, S. A simple sequentially rejective multiple test procedure. *Scandinavian Journal of Statistics* **6,** 65–70 (1979).
14. Benjamini, Y. & Hochberg, Y. Controlling the false discovery rate: a practical and powerful approach to multiple testing. *Journal of the Royal Statistical Society: Series B* **57,** 289–300 (1995).
15. Benjamini, Y. & Yekutieli, D. The control of the false discovery rate in multiple testing under dependency. *The Annals of Statistics* **29,** 1165–1188 (2001).
16. Wright, S. P. Adjusted $P$-values for simultaneous inference. *Biometrics* **48,** 1005–1013 (1992).

17. Reiner, A., Yekutieli, D. & Benjamini, Y. Identifying differentially expressed genes using false discovery rate controlling procedures. *Bioinformatics* **19,** 368–375 (2003).

18. Rowe, C., Harris, J. M. & Roberts, S. C. Corrigendum: Sporting contests: Seeing red? Putting sportswear in context. *Nature* **441,** E3 (2005).

19. Rowe, C., Harris, J. M. & Roberts, S. C. Sporting contests: Seeing red? Putting sportswear in context. *Nature* **437,** E10–E10 (2005).

20. Bradley, R. & Terry, M. Rank analysis of incomplete block designs. I. The method of paired comparisons. *Biometrika* **39,** 324–345 (1952).

21. Seife, C. *Proofiness: how you're being fooled by the numbers* (Penguin Publishing Group, 2010).

22. Hagemann, N., Strauss, B. & Leißing, J. When the referee sees red... *Psychological Science* **19,** 769–771 (2008).

23. Wiedemann, D., Barton, R. A. & Hill, R. A. in *Applied evolutionary psychology* (ed Roberts, S. C.) 290–307 (Oxford University Press, Oxford, 2012).

24. Maier, M. A., Hill, R. A., Elliot, A. J. & Barton, R. A. in *Handbook of color psychology* (eds Elliot, A. J., Fairchild, M. D. & Franklin, A.) 568–584 (Cambridge University Press, Cambridge, 2015).

25. Elliot, A. J. & Maier, M. A. Color psychology: effects of perceiving color on psychological functioning in humans. *Annual Review of Psychology* **65,** 95–120 (2014).

#### Acknowledgments

We thank Tony Black of USA Wrestling, Jeongkang Seo of the World Taekwondo Federation (WTF), Sébastien Gillot of the International Boxing Association (AIBA), "Iceman" John Scully, Janusz Majcher, Andrew Baldwin, Justin Rao, for helpful conversations about Olympic sports. Adam Kenny and Sears Merritt provided research assistance. Mirta Galesic and Michael Lachmann provided helpful feedback on the manuscript.

### Session information

- R version 3.4.4 (2018-03-15), `x86_64-pc-linux-gnu`
- Running under: `Ubuntu 18.04.5 LTS`
- Matrix products: default
- BLAS: `/usr/lib/x86_64-linux-gnu/blas/libblas.so.3.7.1`
- LAPACK: `/usr/lib/x86_64-linux-gnu/lapack/liblapack.so.3.7.1`
- Base packages: base, datasets, graphics, grDevices, stats, utils
- Other packages: dplyr 0.5.0, english 1.1-2, Formula 1.2-2, ggplot2 2.2.1, Hmisc 4.0-3, knitr 1.15.1, lattice 0.20-38, survival 2.44-1.1, xtable 1.8-2
- Loaded via a namespace (and not attached): acepack 1.4.1, assertthat 0.2.0, backports 1.1.0, base64enc 0.1-3, checkmate 1.8.3, cluster 2.0.7-1, colorspace 1.3-2, compiler 3.4.4, data.table 1.10.4, DBI 0.6-1, digest 0.6.12, evaluate 0.10, foreign 0.8-70, grid 3.4.4, gridExtra 2.2.1, gtable 0.2.0, highr 0.6, htmlTable 1.9, htmltools 0.3.6, htmlwidgets 0.9, latticeExtra 0.6-28, lazyeval 0.2.0, magrittr 1.5, Matrix 1.2-14, methods 3.4.4, munsell 0.4.3, nnet 7.3-12, plyr 1.8.4, R6 2.2.0, RColorBrewer 1.1-2, Rcpp 0.12.10, rpart 4.1-15, scales 0.4.1, splines 3.4.4, stringi 1.1.5, stringr 1.2.0, tibble 1.3.0, tools 3.4.4
